# Trajectory inference from single-cell genomics data with a process time model

**DOI:** 10.1101/2024.01.26.577510

**Authors:** Meichen Fang, Gennady Gorin, Lior Pachter

**Author notes:** Address correspondence to Lior Pachter.

## Abstract

Single-cell transcriptomics experiments provide gene expression snapshots of heterogeneous cell populations across cell states. These snapshots have been used to infer trajectories and dynamic information even without intensive, time-series data by ordering cells according to gene expression similarity. However, while single-cell snapshots sometimes offer valuable insights into dynamic processes, current methods for ordering cells are limited by descriptive notions of “pseudotime” that lack intrinsic physical meaning. Instead of pseudotime, we propose inference of “process time” via a principled modeling approach to formulating trajectories and inferring latent variables corresponding to timing of cells subject to a biophysical process. Our implementation of this approach, called Chronocell, provides a biophysical formulation of trajectories built on cell state transitions. The Chronocell model is identifiable, making parameter inference meaningful. Furthermore, Chronocell can interpolate between trajectory inference, when cell states lie on a continuum, and clustering, when cells cluster into discrete states. By using a variety of datasets ranging from cluster-like to continuous, we show that Chronocell enables us to assess the suitability of datasets and reveals distinct cellular distributions along process time that are consistent with biological process times. We also compare our parameter estimates of degradation rates to those derived from metabolic labeling datasets, thereby showcasing the biophysical utility of Chronocell. Nevertheless, based on performance characterization on simulations, we find that process time inference can be challenging, highlighting the importance of dataset quality and careful model assessment.

## 1 Introduction

Single-cell RNA sequencing (scRNA-seq) has provided unprecedented insights into biological dynamical processes in which cells display a continuous spectrum of states that go beyond the confines of discrete cell types [1]. Cells appear to be inherently desynchronized in cellular processes and scRNA-seq can potentially capture cells at different positions over the process even if samples are collected at only one time point. The concept of pseudotime has been developed to describe the position of a cell along the underlying process [2], and trajectory inference (or pseudotemporal ordering) methods aim to solve the inverse problem of inferring the latent pseudotime variable from scRNA-seq data. In light of this this concept, hundreds of methods have been developed [3, 4, 5, 6, 7, 8, 9, 10, 11, 12, 13]. However, with a few exceptions that explicitly model gene expression dynamics [11, 12, 13], trajectory inference methods mostly treat pseudotime as a descriptive concept relying on more or less arbitrary distance metrics in gene expression space. Specifically, there is no well-defined, agreed-upon meaning underlying the notion of pseudotime, and its interpretation is primarily accomplished through qualitative visuals and low dimensional embeddings.

While a descriptive approach can be powerful in exploratory data analysis, the absence of a well-posed definition for a trajectory renders model interpretation and assessment challenging, even conceptually. Firstly, assessing the credibility of results is hard, as fitting can be performed on any dataset and we have limited metrics and ground truth available to gauge the fit quality. Secondly, the interpretation of the inferred trajectory is not straightforward, and downstream analysis based on pseudotime is employed to understand the underlying gene dynamics. However, this need for following analysis gives rise to the problem of circularity (see Sec S1), which becomes evident in the context of an inflated false positive rate in the problem of detecting differentially expressed (DE) genes along pseudotime [14]. To illustrate these two points, we applied the procedure of trajectory inference and DE analysis on simulations generated from four clusters and was able to “discover” superficially plausible dynamics (Fig 2). As naive an example as it is, it reflects the fact that we do not have a reliable way to determine the validity of trajectory inference results. Conversely, adopting a model-based approach has the potential to mitigate this problem. With a clearly defined model of gene expression along a trajectory, the interpretation of parameters and the characterization of errors becomes more straightforward. In addition, the specific question of interest like finding DE genes can be incorporated directly into the formulation of the model, rendering ad hoc analysis unnecessary.

Recently, equipped with a kinetic model of RNA dynamics, RNA velocity has emerged as another powerful concept to provide complementary information about dynamic processes [15]. By distinguishing unspliced and spliced mRNA counts as derived from unique molecular identifiers (UMIs) and fitting gene wise parameters under a on-off model of transcription, it is able to predict the direction of future spliced counts changes. Although a time-dependent gene expression model was explicitly defined in RNA velocity, the time did not have any associated interpretation. Moreover, earlier methods often modeled genes separately with gene-wise times and fit these models after applying a series of ad hoc transformations to count data, which added excessive flexibility and hindered a clear interpretation of the time. As the velocities of different genes had non-comparable scales, they needed to be combined heuristically in a lower-dimensional space to calculate a velocity for a cell [16]. A natural extension is to integrate the cell-wise time of trajectory inference with the mRNA dynamical model of RNA velocity, which a few methods have successfully implemented with different underlying transcription models [17, 18, 19]. Moreover, the recent *VeloCycle* developed an RNA velocity model for the cell cycle that models unspliced and spliced counts dynamics directly with harmonic functions [20].

However, despite the implicit pseudotime modeling performed by some of the RNA velocity methods, there remain many challenges in attaching a physical meaning to pseudotime. Do the parameters of the trajectory model have underlying biophysical interpretations? How can we guarantee that our inferences align with our intended objectives? Are the assumptions of trajectory models satisfied to maintain the consistency of our inference? For instance, the application of trajectory inference or RNA velocity methods relies on the assumption of continuous dynamics in the data, which is not examined retrospectively. Though some heuristic scores are purposed to differentiate between cluster-like data and trajectory-like data [21], there is no principled approach to determine whether the data is sufficiently dynamical and whether a cluster or trajectory model is more appropriate, and the decision of applying trajectory analysis often hinges on prior knowledge and assumptions about the data.

In summary, to attach real meaning to “pseudotime” requires more than just a definition of cell-wise pseudotime. It necessitates a principled approach to statistics to ensure a meaningful inference, which is still lacking in the field of trajectory inference. Meticulous model assessment is required to ensure its relevance to the underlying biological processes and the reliability of results, which includes examining the identifiability of the model, characterizing performance to identify both ideal and failure scenarios, and establishing proper metrics for result falsification. Then pseudotime starts to have a physical meaning, which we suggest defining as “process time” to underscore its interpretation with respect to a specific cellular process.

Therefore, we build such a model and infer “process time” in a principled way with Chronocell. To strike a balance between expressiveness and identifiability, we proposed a trajectory model built on cell states. On the one hand, we incorporated different cell states so that our trajectory model is expressive enough to capture the observation that cells are generally assumed to transition through various states during development. On the other hand, we assume a constant transcription rate for each state to keep the model as simple as possible. By introducing simplifying assumptions in transcription and sequencing model, we ensure model identifiability. We consider the influence of technical aspects and directly incorporate them into the distribution of counts, which eliminates the necessity for unjustified heuristic preprocessing steps that lead to unclear interpretation and biased results even in the large data and no noise limit [16]. We undertook simulations to characterize estimation accuracy in the different parameter regimes to identify ideal and failure scenarios. Using simulations for which ground truth is known allowed us to characterize how large uncertainty and inconsistent parameter values serve as good indicators of potential unreliability in failure scenarios, which can be assessed even when ground truth is unavailable. Finally, we applied Chronocell to biological datasets. We assessed its appropriateness on different datasets and identified unsuitable ones. For suitable datasets, Chronocell revealed distinct cellular distributions over process time and yielded mRNA degradation rate estimates congruous with those obtained from mRNA metabolic labeling.

## 2 Results

### 2.1 A trajectory model generalizing cellular states

We begin by defining the trajectory as a dynamical process underlying all cells, with potentially different lineages/branches within this process. Cells are then assumed to be sampled from various, unobserved, time points along this process. Thus, the latent variable *z* = (*t, l*) is introduced to account for such heterogeneity due to process time (*t*) and lineage (*l*), and *z* follows a sampling distribution determined by the specific biological system and experimental conditions. Consequently, the probability distribution of the data we obtain is a mixture of cells over time and lineage,

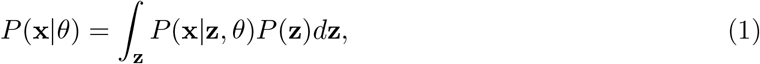

where **x** is the data and θ is the set of parameters that define the trajectory. This is the common framework of trajectory inference, and developing a trajectory model requires defining the gene dynamics along the lineage and the process time *P* (**x**|**z**, θ), as well as the sampling distribution of cells over the process *P* (**z**).

To define the dynamical process, we first state our transcription model. We consider only transcription, splicing and degradation reactions in cells, and assume only transcription rates are time-dependent (Fig 1 Gene expression):

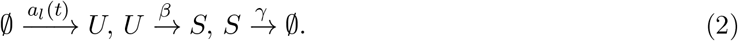

where *a*_*l*_(*t*), *β* and *γ* are the transcription, splicing, and degradation rates for lineage l. The chemical master equation describing the evolving distribution of the above biochemical reaction network has an analytical solution [22]. If we assume the initial distribution to be Poisson, the solution remains Poisson with the means (*λ*_*u*_ and *λ*_*s*_) of *U* and *S* evolving according to the following ordinary differential equations (ODEs) of RNA velocity:

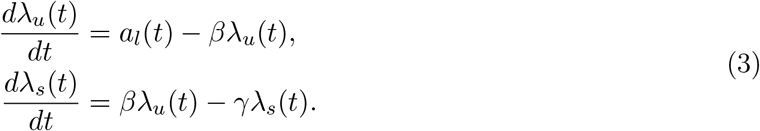

**Figure 1.**
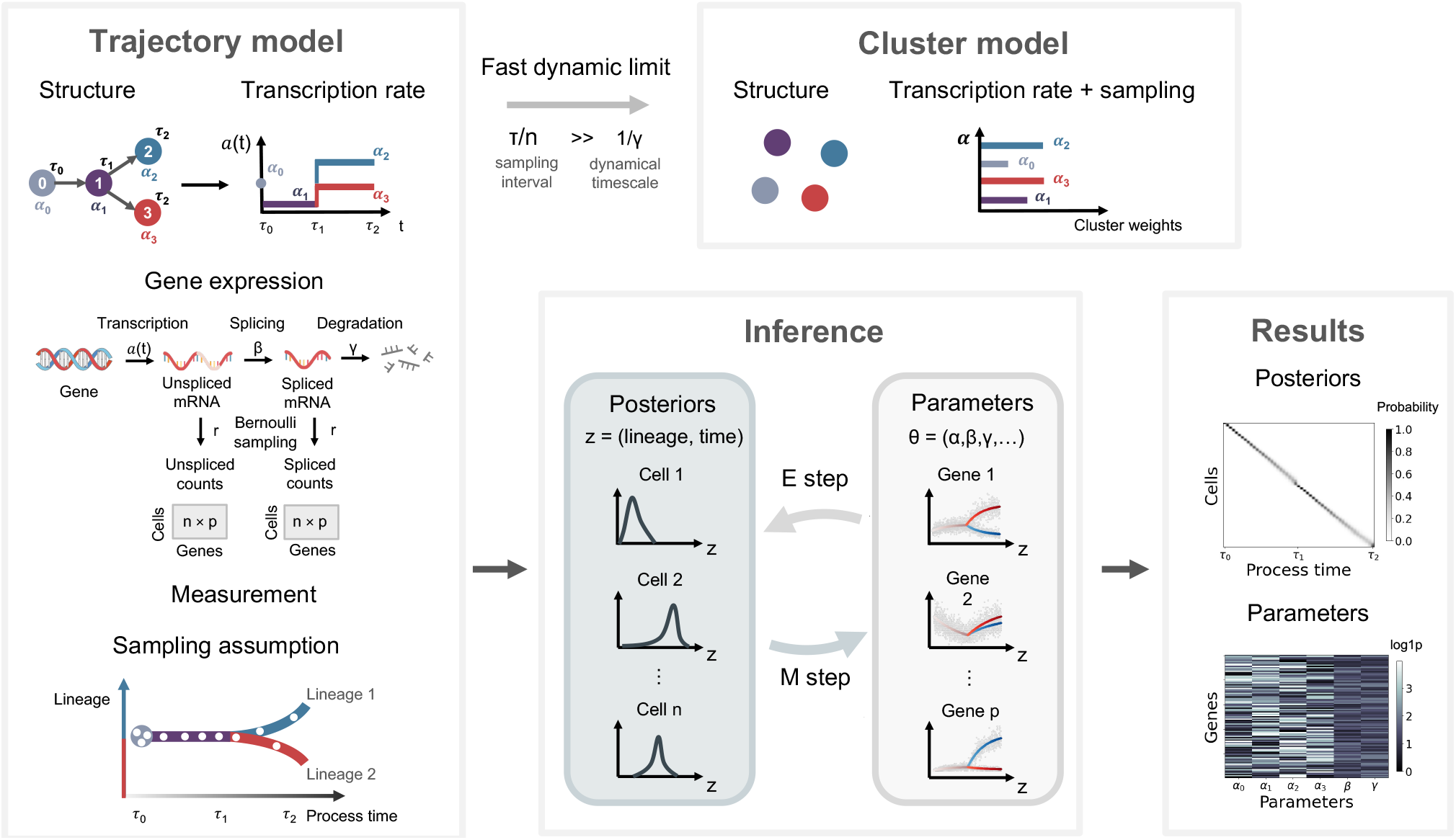
Chronocell model and inference. The **trajectory model** of Chronocell is associated with a trajectory *structure* that specifies the states each lineage goes through. *Transcription rate*s are modeled as a piecewise deterministic function for each lineage. Each state *s* is associated with a transcription rate α_*s*_ and exit time *τ*_*k*_. For synchronized model, all genes share the same *τ*_*k*_, while for desynchronized model, each gene can have a different *τ*_*k*_. Transcription, splicing and degradation reactions are considered in the *gene expression* model. Binomial sampling with cell-wise read depth *r* is used to model the *measurement* of mRNA in scRNA-seq. The *sampling assumption* specifies the distribution of cells across lineages and process time: the distribution over process time (0,1] is fixed to be uniform, and the point mass on time 0 and weights of lineages are treated as parameters determined through fitting. A trajectory model degenerates into a **cluster model** (Poisson mixtures model) in the fast dynamic limit where the dynamical timescale is much smaller that the cell sampling timescale. The mixture weights of clusters correspond to the product of the length of corresponding time interval (or mass on time 0) and weights of lineages of the sampling assumption. An EM algorithm is used for **inference** of trajectory models, and *parameters* and *posterior* distributions are fitted iteratively, which are the direct **results** of a trajectory model.

**Figure 2.**
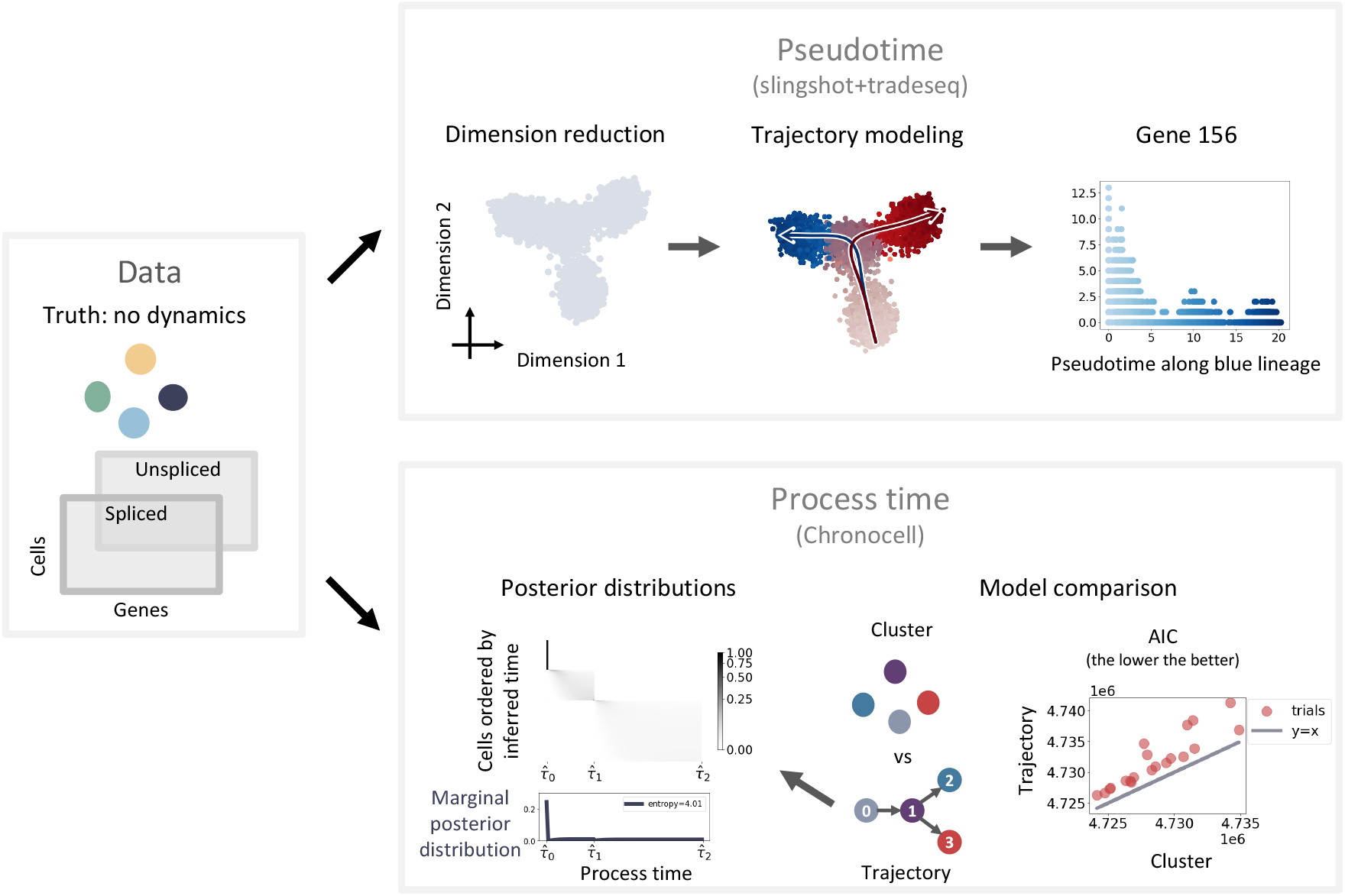
Principled process time inference enabled by Chronocell. **Data** are simulated from 4 discrete clusters (4 Poisson mixtures). As an example of **Pseudotime**, trajectory in lower dimensional space was constructed with Slingshot [9]. Differential genes along pseudotime were selected with tradeSeq [23], with one gene plotted along the blue lineage. An example of **Process time** where we use Chronocell on the same data. Posterior distributions of cells on process time grids are plotted in gray scale, with the average posterior distribution over cells below. For model comparison, AIC scores of Poisson mixtures model and trajectory model of 20 samples from the same parameter set are compared.

The modeling of transcription rate *a*_*l*_(*t*) is motivated by the common abstraction of cellular differentiation as cell state transitions: each lineage is abstracted as a series of switches in cellular states over time (Fig 1 Structure). We assume transcription is constitutive and neglect transcriptional bursting, and introduce one transcription rate per gene for each cellular state (α). Switching is assumed to be instantaneous and occurs at an unknown but fixed time (*τ*) with the first switch to occur at *τ*_0_. Without loss of generality, we consider the entire process to start at time 0 and have a time length of 1. Consequently, the transcription rate function *a*_*l*_(*t*) of lineage l is simplified as piecewise constant functions of the process time over [0, 1] (Fig 1 Transcription rate). This piecewise constant function is defined in the limiting regime where transcriptional state switching (such as expression of master regulatory factors and changes of chromatin state) precedes gene expression and has a much faster time scale. It also directly reduces to discrete cell clusters in the fast dynamic limit, i.e., when dynamical timescale (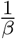 and 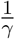) is much smaller than sampling intervals, for example, the total time length divided by cell number, *n*, under a uniform sampling distribution (Fig 1 Cluster model). This connection to cluster models enables us to interpolate between discrete cell states and continuous dynamics.

The simple form of the transcription rate function and the lack of bursting lead to a tractable model and facilitates inference and analysis. In fact, it affords explicit solutions for the distribution and for its derivatives with respect to parameters. Ideally, gene regulatory networks involved in cell differentiation would be modeled, but with current data types it is difficult to include gene interactions and to model transcription rates as functions of other genes with accuracy. Thus, we assume that the dynamics of different genes are independent and that all correlations are absorbed into the shared latent process time. We also assume all genes are roughly synchronized in the basic model, i.e. *τ* is the same for all genes, but the choice of desynchronized switching (different *τ* for each gene) is also provided in the desynchronized model. In summary, our trajectory model is suitable for capturing the coordinated global gene expression changes instead of the detailed gene dynamics.

After deriving an explicit distribution of in vivo counts, we turn to the sequencing model. We assume simple binomial sampling of each molecule, and that the average binomial sampling probability, i.e., read depth, varies between cells but remains the same for all molecules in one cell (Fig 1 Measurement). Then, in vitro counts remain Poisson but with means adjusted by read depth. Instead of using normalized data, we estimated read depth using the total UMI counts of near-Poissonian genes that are not used for the inference, and incorporated it into the count distribution. Therefore, we arrive at an analytical form of the conditional probabilistic distributions *P* (**x**|**z**, θ) of counts from a dynamic process by specifying the trajectory structure, gene expression model, transcription rate functions, and scRNA-seq measurement model.

The remaining part of specifying the sampling distribution *P* (**z**) is crucial, because it breaks the scale invariance of parameters and ensures the identifiability of the model: multiplying the transcription, splicing and degradation rates by the same constant leads to the same marginal likelihood if the sampling distribution can be changed. Ideally, the specification of the sampling distribution depends on our knowledge of the studied biological system and the experimental design. For example, for stem cells that constantly divide and differentiate, we may assume a uniform sampling distribution over (0, 1] with a point mass on time 0, where the point mass represents the fraction of time that cells spend in the initial state. On the other hand, for time series data, we may assume the sampling distribution of cells were centered around their captured time points with some variances. However, in practice, since there is no obvious principled way to determine such distributions, we just assume a uniform sampling distribution over process times (*t* > 0) by default, but the weight on time point 0 and different lineages can be identifiable and be updated in the inference.

A trajectory is thus defined with the above dynamical process and sampling distributions. With the parameterized form of the probabilistic distribution of counts, we can employ the Expectation-Maximization (EM) algorithm to estimate model parameters and posterior distributions of process times and cell lineages by maximizing Evidence Lower Bound (ELBO) with either warm start or random initialization (Sec 4.2). The synchronized model was used by default, where all genes share the same *τ*. The fitting of desynchronized model also starts from the result of the synchronized model. For desynchronized models, we also introduce a penalty term proportional to the squared difference between gene-wise *τ*_*k*_ and the global *τ*_*k*_ to encourage synchronization, and the coefficient for this penalty term can be specified as a parameter (0 by default). Both models are identifiable and parameters can be recovered accurately (Fig S1 and Fig S2).

By having an explicitly parameterized distribution of raw counts, we can easily interpret results and systematically assess the model under a more principled framework. Since parameters all have biophysical meanings, we can not only directly interpret them but also validate their accuracy by comparing them to orthogonal experiments that measure the same parameters. Furthermore, we can directly select DE genes by fold changes in transcription rates across states after filtering genes by goodness of fit. The performance can be quantified through parameter errors, aiding in the identification of both confident and uncertain scenarios. As for false positives arising from clustered data, we can evaluate when a trajectory model is no longer appropriate by comparing it to the cluster model using standard model selection methods like AIC (Fig 2 Process time). Importantly, the posterior distributions and multiple minima of AIC scores can hint at the lack of continuity (Fig 2 Process time), serving as a retrospective metric when model selection methods are compromised by unaccounted noise.

### 2.2 Demonstrating performance of Chronocell on ground-truth simulations

We first demonstrate the inference and evaluate its performance in the perfect scenario, where simulations are generated strictly under our trajectory model without any unaccounted noise. We used a two-lineage trajectory and uniformly sampled 10,000 cells over lineages and process times with biological plausible parameters extracted from literature [24] (Sec 4.4). 100 out of 200 genes are variable (Fig. 3a). We fit under a slightly wrong assumption of the trajectory structures and assumed all genes are variable (Fig. 3b). For random initialization, we initialized randomly 100 times and picked the one with highest ELBO score (Fig. 3c). We also fit with warm start by initiating the fitting process from correct cell clusters grouped by true time and lineages. Both types of initialization were able to converge to the ELBO with true parameters, and random initialization yielded a slightly higher ELBO compared to warm start, with a negligible difference (Fig. 3d). For the following analysis, we used the fitting results of random initialization. The posterior distributions correctly recapitulate the time and lineages of cells, with a root mean squared error (RMSE) around 0.05 for the posterior mean of process time in comparison to the true time (Fig. 3e). This means that the error in time of a cell is around 5% of the total time length of the trajectory in average. However, the error is not completely uniform: as cells in the second interval are closer to the steady state, they have more spread posteriors and larger errors. Since the posteriors of each cell are accurate, the posteriors averaged over cells also resemble the empirical distribution of the true time (Fig. 3f).

**Figure 3.**
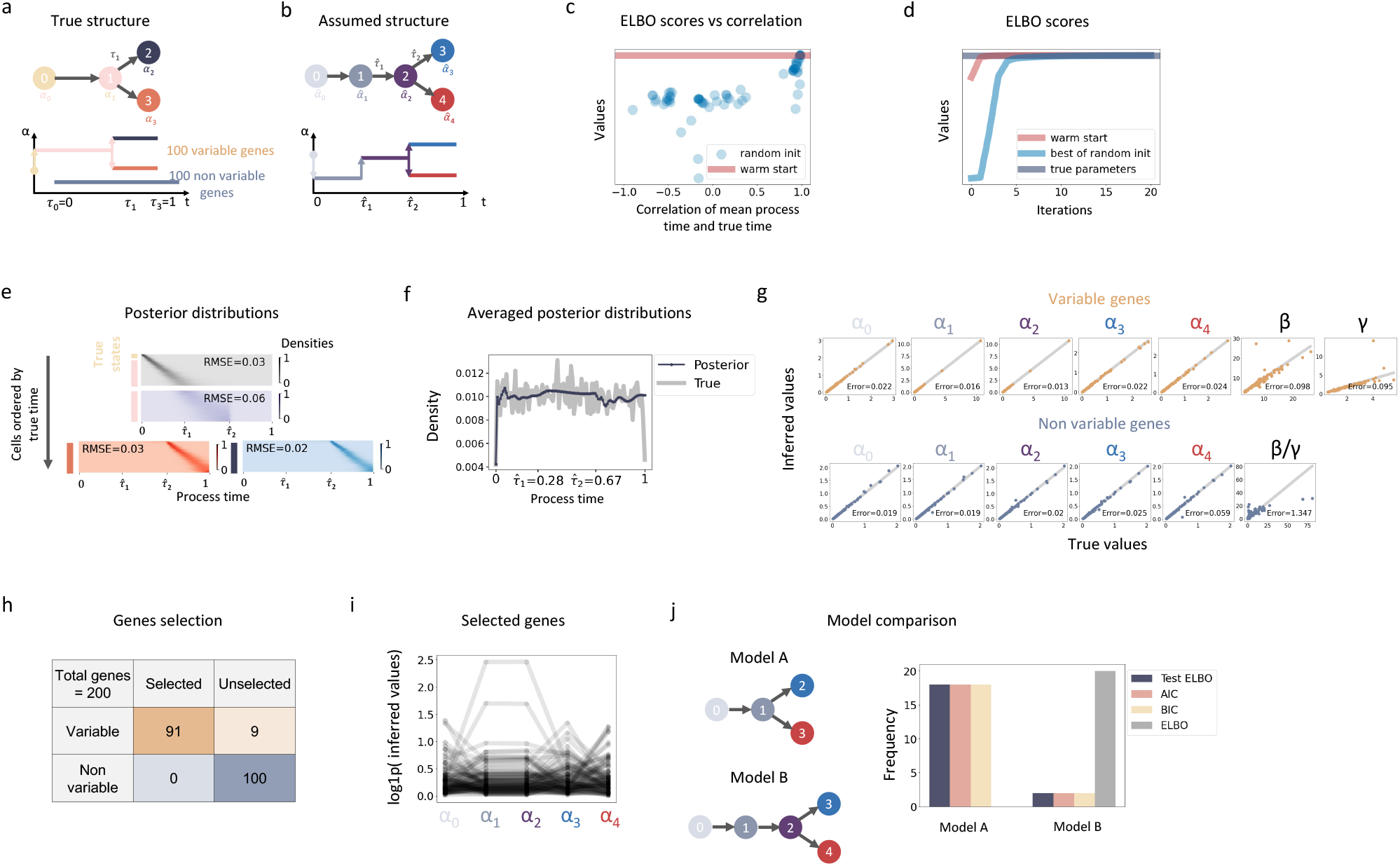
Demonstration of inference on simulation. **a** The ground truth trajectory structure. Cells jump to the next state (2) from starting state (1) at *τ*_0_ = 0, and then bifurcate into two lineages with different ending states (3 and 4) at *τ*_1_. The process ends at *τ*_2_ = 1. Out of 200 total genes, 100 genes are non variable with the same distributions along time. **b** The assumed structure that does not know the first two states are supposed to be merged into one. All genes are assumed to vary along time. **c** The ELBO scores of 100 random initializations (blue dots) compared to those of warm start (red line). The x axis is the Pearson’s correlation between the mean process time of each random initialization and the true time. **d** ELBO scores over fitting iterations of both warm start (red line) and the best random initialization (blue line), with the ELBO calculated with true parameters (gray line) as reference. **e** Heatmaps of inferred posterior distributions. x axis is time grids, and y axis is cells aligned by their true times with true transcription states on the left. The intensity of color indicates the weights of posterior distributions of cells on the grids. Heatmap of cells from *τ*_0_ to *τ*_1_ use a gray color palette. Heatmap of cells from *τ*_1_ to *τ*_2_ use purple color palette. Heatmap of cells from *τ*_2_ to *τ*_3_ of first lineage use blue, and those of second lineage use red. RMSE stands for root mean square errors. **f** The averaged posterior distributions across cells (dark blue) and true empirical distribution (gray) of process time. **g** Inferred parameters values compared to true values. For non variable genes, only 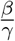 are identifiable and compared. Error is mean normalized error as described in the text, and the mean is computed across genes. **h** The confusion matrix for gene selection. **i** α values of selected genes over states. **j** Two models and the distribution of the chosen one by (train) ELBO, AIC, BIC and test ELBO, calculated on 20 samples each with a different set of parameter.

The parameters are also recovered accurately (Fig. 3g). For variable genes, all parameters are identifiable, while for non-variable genes, only *α* and the ratio of *β* to *γ* are identifiable. We noticed a trend that the absolute errors of α scale with the square root of true values, while for *β* and *γ*, they scale with the true values (Fig. S3a). This trend also appeared in other simulations (Fig. S1b,c). Therefore, to calculate the errors of parameters, we divide the absolute error of α by the square root of true values and *β, γ* by true values, so that errors of different genes are more comparable. We refer to them as normalized errors in the text. The parameter α tends to be estimated with higher accuracy compared to *β* and *γ* (Fig. 3g). This aligns with intuition because, although parameters are identifiable in our trajectory model, the Evidence Lower Bound (ELBO) is nevertheless insensitive to proportional changes in *β* and *γ*, which have minimal impact on the phase portrait in the unspliced and spliced space of each gene. To confirm this, we used the true model to calculate the Fisher information matrix and the smallest eigenvalues of each gene (Fig. S3b). The corresponding eigenvectors describe the flattest direction of the ELBO. To validate that the flattest direction primarily lies in the *β* and *γ* parameters, we add the corresponding eigenvectors to the true parameters after normalizing it by the square root of eigenvalues, and calculate the new ELBO with the modified parameters. Indeed the resultant changes in ELBO are indeed small and *β* and *γ* values were mainly varied (Fig. S3c), which confirms that the *β* and *γ* are harder to estimate accurately (Fig. 3g).

To select genes whose dynamics are well-fit by the model, we evaluate their goodness of fit by comparing the gene-wise likelihood with that of clustering models (See Sec 4.3.2). We adopt this relative likelihood criterion because absolute likelihoods of different genes are not directly comparable and clusters model serves as a natural reference point for comparison to filter genes without continuous dynamics. All 91 genes selected belong to the variable class (Fig. 3h). The similar transcription rates of states 1 and 2 of selected genes would also suggest us that those two states could be merged into a single one, if we didn’t know the true structure (Fig. 3i). Therefore, we applied model selection methods to compare the ground truth model and the assumed model. We generate another 20 parameter sets, fit under both models, and compute in-sample ELBO scores (ELBO), AIC, BIC, and out-of-sample ELBO scores (test ELBO). The ELBO is always better in model B due to overfitting. All of the last three metrics (AIC, BIC and test ELBO) favor the true model most of the time (90%) (Fig. 3j). However, we want to emphasize that model selection worked because simulations were strictly generated under our trajectory model. In reality, if true transcription rates are far away from piece-wise constant functions, the resulting deterministic noise can introduce bias in AIC/BIC and cross-validation, leading them to prefer more complex models [25].

### 2.3 Identifying failure scenarios reveals the fragility of inference

Given the accuracy on perfect data, we sought to identify potential failure points that can undermine the accuracy of the inference and characterize their impact. In addition to obvious factors like insufficient numbers of cells or genes as well as low read counts (Fig S4), we tested the effects of noise on inference, as unaccounted noise is prevalent in scRNA-seq datasets. The first type of noise is cell-wise read depth (or cell size) that influences all genes similarly. The inference is sensitive to such noise: when read depth variance is not correctly accounted for in the fitting, inference results may be highly inaccurate when its squared coefficient of variation (CV^2^) exceeds 0.1 (Fig. S5a). As the CV^2^ typically observed in real datasets exceeds 0.1 (Fig S10), an accurate estimate of cellwise read depth is critical. Considering this fact, instead of simply using total counts as normalization factors, we use normalized covariance between genes to decompose extrinsic noise (influencing all genes) and intrinsic noise (gene specific). We estimate the read depth CV^2^ across cells from the normalized covariance between genes. Subsequently based on the read depth CV^2^, we subtract the extrinsic variance caused by the read depth from total variance and identify Poissonian genes whose remaining variances (intrinsic variances) are close to their means (variance < 1.2 mean). We then estimate cell-wise read depth using the sum of those Poissonian genes. Different sets of genes are used for estimating the CV^2^ of read depth and fitting trajectories. In reality, these read depth estimates correlate well with total counts number for most datasets, with one interesting exception (Fig S10).

On the other hand, gene specific noise seems to have less impact on inference. We added gene-wise gamma noise in simulation which generates negative binomial distributions in steady states and approximates bursting noise. Parameter errors gradually increase as CV^2^ of Gamma noise increases but remain reasonably small even with a CV^2^ of 1 (Fig. S5b). However, while it may not completely undermine the fitting process, it can lead to failures in model selection. When repeating the procedures in Fig 2 and Fig 3h after introducing Gamma noise, the AIC/BIC and cross-validation metrics favor the wrong models that were more complex than the true one (Fig. S5c and d). This highlights the importance of providing reasonable trajectory structure to the fitting based on prior knowledge, and exploring alternative methods for quality control against false positives caused by clusters.

Another probable factor in real datasets that can lead to suboptimal results is is insufficient dynamics. Intuitively, for a dynamical model to be appropriately fitted, the data must capture a significant amount of transient dynamics to recapitulate the evolving processes over time, without which the data start to resemble discrete clusters. Such cases can occur in at least two possible scenarios: fast timescales and concentrated sampling distributions, both of which result in clusters in the extreme. Therefore, false positives caused by clusters are naturally included as a component of identifying unreliable results.

The first situation, fast timescales, arises when mRNA half-lives are significantly shorter than the timescale of biological processes, thus cells are mostly near steady state and provide little information about the intermediate dynamics. By setting the length of the time interval to unity and increasing *β* and *γ*, which is equivalent to increasing processes timescale with unchanged mRNA half-lives, we found that ideally, the mean value of *γ* should not fall out of the range of 1 to 10, and the mean ratio of *γ* to *β* over genes should not be too small (Fig. S6). Hence, if the average half-life of spliced mRNA is approximately 30 minutes [24], snapshot data sampled from processes involving steps exceeding 10 hours are no longer suitable.

Furthermore, given an appropriate timescale, the sampling distribution still has to cover the region where the transient dynamics occur. Imagine, for example, that all cells are from one unknown time point; there is no way for a trajectory to be inferred. Instead, a cluster should be used to fit the data. Thus, it is crucial to determine the minimum level of uniformity required in the sampling distribution and verify whether this requirement is met. By gradually changing sampling distributions from a uniform distribution to a Gaussian with a random mean, we generate datasets with sampling distributions that exhibit decreasing levels of uniformity, which was quantified using entropy (Fig. S7a). As the sampling distribution deviates from uniformity, the errors of process time and parameters quickly increase as expected (Fig. S7b). In fact, even when the sampling distribution is not far away from a uniform one with entropy 4, the errors are big enough that the results are completely wrong (Fig. S7b). Further, even using the true sampling distribution as a prior for a warm start with correct initialized time cannot mitigate the lack of dynamics: the errors in parameter estimations remained substantial (Fig. S7c). Therefore, sufficiently transient dynamics is an inherent requirement for trajectory inference even when perfect prior information is provided.

In summary, a suitable dataset for fitting process time needs to contain enough dynamic information as well as limited noise. This stringent requirement for a successful trajectory inference highlights the need for suitable datasets and careful model assessment. Straightforwardly, we could make a consistency check by verifying if the remaining noise, parameter values, and uniformity of the average posterior distribution fall into the appropriate range determined in simulations. However, when a result is incorrect, it may not necessarily exhibit large unexplained noise, high splicing/degradation rates, or a concentrated cellular distribution over process time, as it could inadvertently fit undesired patterns and output plausible results. Thus, inconsistency is a sufficient but not necessary indicator of unreliable results.

It turns out that the large uncertainty of results can be a better indicator. As both noise and limited dynamics tend to diminish or obscure the difference of scores like ELBO between correct and incorrect outcomes, they introduce multiple comparable maxima and make the fitting results unstable. The resultant large uncertainty can be measured by two different approaches. First, as each random initialization outputs a different ELBO/AIC score and cell ordering, we can inspect the distribution of scores with respect to a summary parameter of cell orderings, which describes the global landscape of the score function. Ideal scenarios usually give one distinct maximum of ELBO (or minimum of AIC) at the correct ordering of cells (correlation around one), while failure scenarios usually lead to multiple comparable maxima of ELBO (or minima of AIC) at different cell orderings beside the correct one (Fig. S8b). We use average precision (AP) of the 100 random initializations to quantify the distinctiveness of the correct outcomes and summarize the uncertainty. One random initialization is considered correct if its resultant mean process time correlates well with that of the best one, e.g., a Pearson’s correlation of 0.8 (Sec 4.3.3). Ideal cases lead to AP close to one and low AP indicates instability but not vice versa, which means low AP is a sufficient indicator for instability (Fig. S8b). Second, the uncertainty can also be revealed by standard bootstrap analysis. We generated 100 sets of resampled data and computed the correlation in process time between the original and each resampled set (Sec 4.3.3). In failure scenarios the results of original data and resampled data often differ and correlations are scattered with large variance. While in ideal scenarios, process time estimates of resampled results agree with the original one and correlations are centered around one (Fig. S8c).

### 2.4 Uncovering distinct underlying cellular distributions in process time

We applied Chronocell to a variety of datasets with different anticipated sampling distributions over time. For those datasets, we estimated read depth as described in 4.2.2 (Fig S10), and then filtered genes for fitting based on their means, variances and unspliced to spliced ratios (see Sec 4.5). The trajectory structures are determined based on prior knowledge. Random initialization was always performed and its uncertainty was assessed by both AP and bootstrapping. Warm start was applied as well if cell type annotations were available, which were used to initialize the fitting process. The corresponding clusters (Poisson mixtures) model was fitted with the same read depth and used for comparison. The genes were selected in the same manner as in the simulation. DE genes are selected based on the fold changes.

We first tested our method on the T cells of PBMC (Peripheral Blood Mononuclear Cells) dataset from 10x Genomics, which is typically expected to exhibit a few distinct clusters (Fig S11a). Indeed, though the AIC of trajectory model is lower than clusters model likely due to unaccounted noise, the scores of 100 random initializations display multiple minima: multiple different cell orderings result in similar AIC values (Fig S11b). The low average precision (0.28) of random initializations suggests clusters model would be more suitable for PBMC. Serving as a negative control, it confirms our ability to reject unreliable results on real datasets, even when standard model selection methods are invalidated by incorrect modeling of noise.

The second dataset contains a snapshot collection of glutamatergic neuronal lineage cells in a developing human forebrain (Fig 4a) [15], which is presumed to capture cells along a continuous trajectory. However, the unstable result of random initializations indicates its unreliability, as results with reversed directions yield comparable AIC scores (Fig 4b), resembling simulations that lack dynamics information (Fig S8b). This observation is further supported by examining the cellular posterior distributions obtained through warm start with cell type annotations, where the average posterior distribution reveals that cells are concentrated around starting time *τ*_0_, i.e., the initial state, with a low entropy (Fig 4c). Therefore, both consistency and instability checks suggest this is not a suitable dataset for Chronocell.

**Figure 4.**
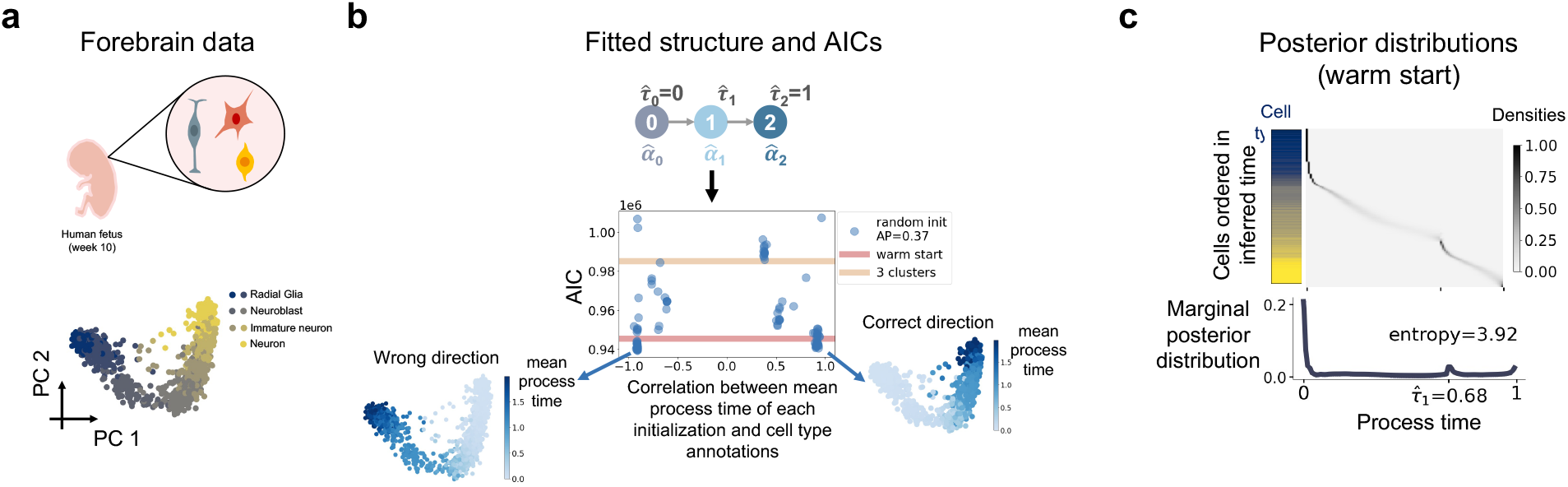
Inference results for Forebrain data. **a** Schematics of Forebrain data and PCA plot of cells colored by cell type annotations. **b** The fit trajectory structure, and AIC scores and mean process time correlations of 100 random initializations (blue dots) compared to those of warm start (red line) as well as 3 clusters (Poisson mixtures) model (yellow line). AP stands for average precision. Inferred mean process time of two initializations with opposite directions are indicated in blue on the same PCA plot as in **a. c** Posterior distributions of process time. Cells were ordered in y axis by their inferred mean process time and the left bar displays their cell types using the colors in **a**. The below histogram shows average posterior distribution averaged over cells. The entropy of the average posterior distribution was calculated using its weights on the 100 discretized time grids.

The third dataset contains erythroid lineage cells during mouse gastrulation collected from multiple time points (Fig 5a) [26]. Biological time availability offers a valuable means to evaluate results by comparing the posterior distributions of cells to their corresponding physical times, which is particularly useful since real snapshot datasets lack a definitive ground truth. The AIC scores of random initializations show a clear minimum at a correct direction that align with cell type annotations (Fig S12a), and the average precision of random initializations is 0.79 which is notably higher than those of PBMC and Forebrain datasets (Fig S11b and Fig 4b). The process times of bootstrap samples are reasonably stable as well (Fig S12a). The fitted dynamics were able to explain most of the variance, leaving the CV^2^ of unexplained noise of most genes under 1 (Fig S12c).

**Figure 5.**
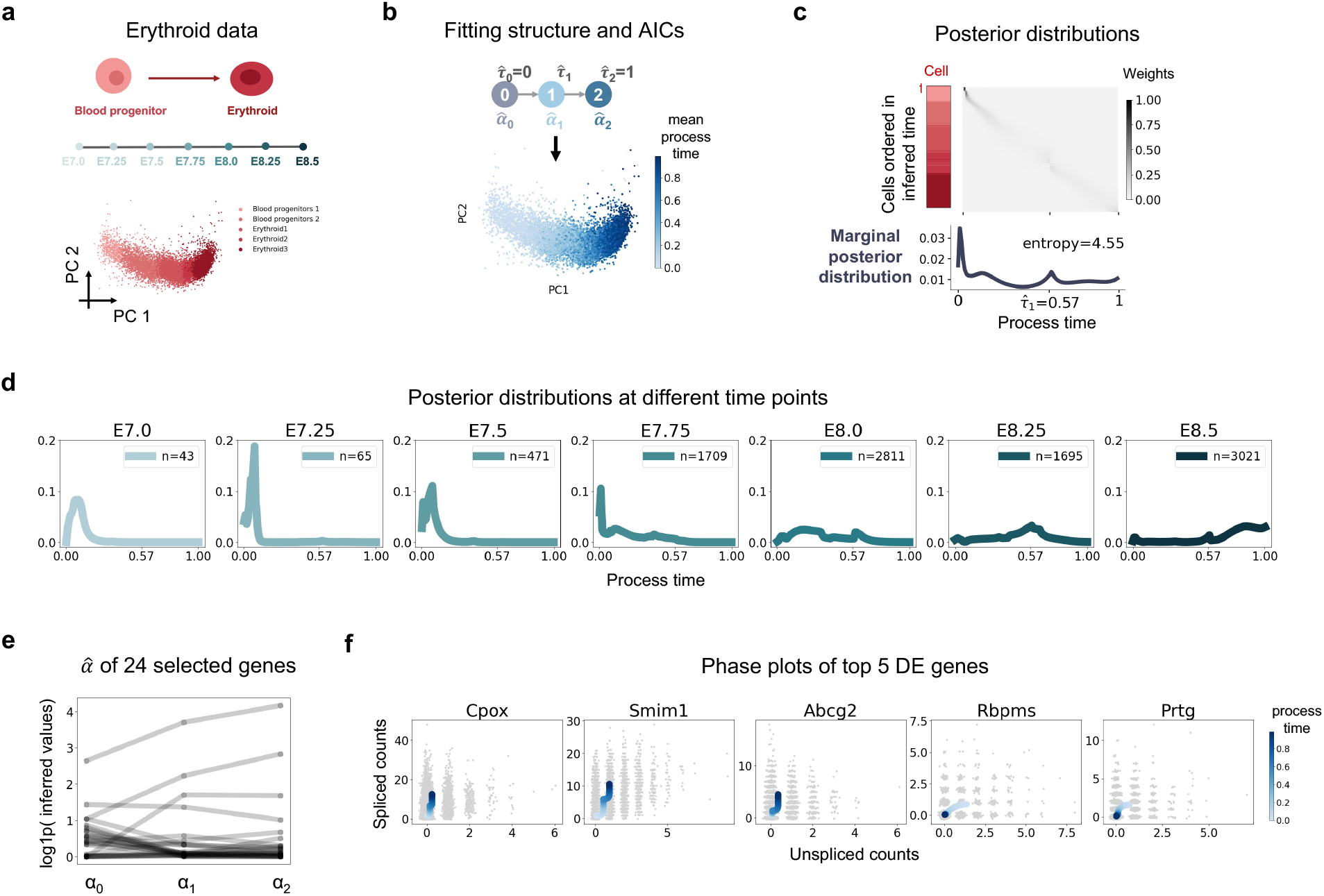
Inference results for Erythroid data. **a** Schematics of Erythroid data and PCA plot of cells colored by cell type annotations. **b** The fitted trajectory structure and inferred mean process time from random initialization indicated in blue on the same PCA plot as in **a. c** Posterior distributions of process time. Cells were ordered in y axis by their inferred mean process time and the left bar displays their cell types using the colors in **a**. The below histogram shows average posterior distribution over cells. The entropy of the average posterior distribution was calculated using its weights on the 100 discretized time grids. **d** Averaged posterior distribution across cells from different experimental time points. n is the number of cells. **e** α values of 24 selected genes over states. **f** Phase plots of top 5 DE genes of 24 selected genes. The x axis is the raw unspliced counts and y axis is the raw spliced counts. The blue curve is the fitted mean of product Poisson distributions of unspliced and spliced counts over process time, and its darkness corresponds to the value of process time.

Therefore, the Erythroid dataset appears to be a suitable dataset for our trajectory model. The best result from random initialization is used for analysis (Fig 5b), and cellular posterior distributions confirm that cells have a broad distribution over process time (Fig 5c). What’s more, the posterior distributions of cells from different time points roughly move forward along the process time in sync with real-time progression (Fig 5d). Although not perfect, it suggests that our trajectory model can successfully capture the correct trend of process time, even when assuming a uniform sampling distribution for all cells.

After applying our gene selection procedure based on relative likelihood and discarding genes with extreme values, we ended up with 24 (49%) genes (Fig 5e). Subsequently for demonstration, we chose the top 5 DE genes (*Cpox, Smim1, Abcg2, Rbpms, Prtg*) with the largest fold change of transcription rates, and plotted their phase portraits (Fig 5f). Interestingly, four of them (*Cpox* [27], *Smim1* [28], *Abcg2* [29], *Rbpms* [30]) were reported to be directly relevant to erythroid development, which illustrates that selecting DE genes based on inferred transcription rates is a straightforward and effective approach.

Based on the results from datasets ranging from clusters to trajectories, it becomes evident that real datasets can display a spectrum of continuity in cellular process time distribution. Therefore, it is critical to assess the quality of inference and verify the requirements for reliable results are indeed met.

### 2.5 Degradation rates estimates agree with metabolic labeling data

In addition to validating the process time, we also sought to validate the inferred parameters, which motivated us to use a metabolic labeling dataset from scEU-seq, comprising human retinal pigment epithelial (RPE1) cells undergoing cell cycles (Fig 6a) [31]. Metabolic labeling of new mRNA allows for the estimation of degradation rates from cells with varying labeling times, enabling a comparison with our parameter estimations.

**Figure 6.**
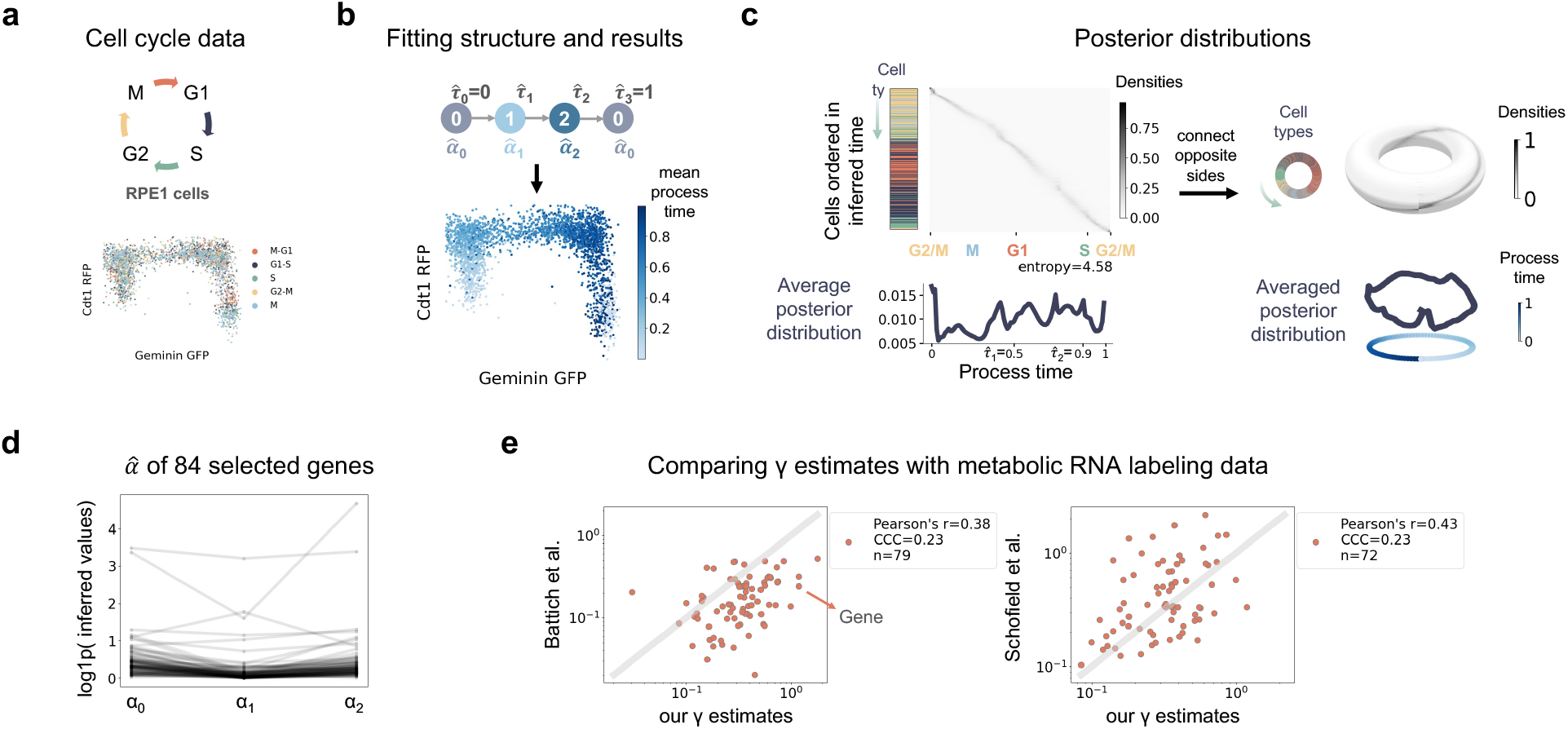
Inference results for cell cycle data. **a** Schematics of cell cycle data and scatter plot of the Geminin-GFP and Cdt1-RFP of RPE1 cells colored by cell type annotations. **b** The fitted trajectory structure and inferred mean process time from random initialization indicated in blue on the same scatter plot as in **a. c** Posterior distributions of process time. Cells were ordered in y axis by their inferred mean process time and the left bar displays their cell types using the colors in **a**. The below histogram shows average posterior distribution averaged over cells. The entropy of the average posterior distribution was calculated using its weights on the 100 discretized time grids. **d** α values of 84 selected genes over states. **e** Comparison of *γ* estimates for 84 selected genes with estimates derived from metabolic RNA labeling data. CCC stands for concordance correlation coefficient. n is the number of genes for which estimates are available in each respective paper.

In the trajectory structure, we specified that the initial and final states were identical. Given our assumption that the initial state is at a steady state, capturing the cyclic nature of the cell cycle poses an additional requirement that cells must also approach a steady state in the last interval as well, which happened to be reasonably satisfied by our results. In line with the expectation that cell cycle is a highly dynamic process, the random initialization provides a relatively stable result: the AIC exhibits a single minimum that aligns with the direction of cell cycle progression (Fig S13a), and bootstrap samples mostly align with the result of the original one (Fig S13b). Starting from the result of random initialization, a desynchronized model was fitted to enhance the accuracy of parameter estimations (Fig S13c). The resulting fit maintains alignment with the cell cycle progression (Fig 6b), and the CV^2^ of the unexplained noise mostly remains below 1 for most of the genes (S13c). The RPE1 dataset was metabolically labeled for different lengths of time but posterior distributions of RPE1 cells with different labeling times do not show significant differences, as a negative control in contrast to the erythroid data (Fig S13e).

The posterior distributions successfully capture the cyclic nature of the data. The cell types ordered by mean process time exhibit a cyclic pattern (Fig 6c); the phase plots of marker genes confirm the cycling dynamics of fitted means of unspliced and spliced counts (Fig S13f); the starting and ending values of fitted means of most of genes match with each other (Fig. S13g). As both axes are cyclic, it is possible to connect the opposite edges and transform the two-dimensional cell-by-time grids posterior distributions into a torus on the surface of which the posterior distributions look like a circle (Fig 6c). Based on cell type annotation, total RNA counts number (Fig S13h), and marker genes dynamics (Fig S13e), we can roughly assign the three intervals to G2/M, G1/S and S/G2 phase, and mitosis happens shortly after *τ*_0_.

After selecting genes by relative likelihood and discarding genes with extreme values, we ended up with 84 (46%) genes (Fig. 6d). We compared the degradation rates of 84 selected genes to those derived by scEU-seq [31] and TimeLapse-seq [32]. For scEU-seq, as we neglected the changes in degradation rate along cell cycle, we used the averaged degradation rates in Fig. S10C of Battich et al. We observed moderate correlations between our degradation rate estimates and the respective estimates from Battich and Schofield (Fig. 6e), and these correlations exhibited a magnitude similar to the correlation between the Battich and Schofield estimates (Fig. S13i), which suggests that the parameters inferred by Chronocell indeed possess a meaningful biophysical interpretation.

## 3 Discussion

We have introduced Chronocell, a method constructed upon a trajectory model featuring biophysically meaningful parameters and a principled approach to fitting and analysis. Counts are directly modeled, estimation accuracy is characterized, and unreliable instances are discerned. We found several requirements regarding dynamics and noise that must be satisfied to obtain reliable results, which makes process time inference challenging and renders retrospective model assessment indispensable. We applied Chronocell to different kinds of real datasets, recognized inapplicable ones by assessing instability, and demonstrated meaningful interpretation of process time and parameters on applicable ones.

Of note, our trajectory model is simplified to balance interpretability and tractability. Building on the concept of cell state transitions, we have assumed piecewise constant transcription rates which describe genes with fast chromatin states switching in saturated regime, but may be unrealistic for many biological systems. Furthermore, we have assumed that splicing and degradation rates remain constant, which although reasonable for many genes, is not accurate for genes with peaked response [24]. We also did not incorporate transcriptional bursting due to the absence of an analytically determined temporal solution, and this leads to a under-dispersed distribution compared to what is typically observed in biological data. Nevertheless, this can be resolved by using numerical solutions [33]. However, even with this simplified model, we have found that accurate inference imposes strict requirements on data. This question is inherently challenging and insufficiently specified due to the existence of latent variables and flexibility of the transcription rates. A more realistic model would have even more stringent requirements.

Since fitting dynamical parameters is challenging from static snapshots, physical time information could offer valuable insights when incorporated into our latent variables model. While the requirements on sampling and noise might potentially be relaxed, they would likely persist to some extent. Thus, intensely sampled time series datasets derived from well-defined cellular processes would be the ideal choice for trajectory inference. However, it remains to address the key question of how physical time should be translated into sampling distribution assumptions. Should we fix the process time to physical time completely, i.e., a delta distribution? Or should we assume heterogeneity within one time point? If so, what kind of distribution should we use? There is no straightforward answer, and assuming different distributions (delta, exponential, uniform) can have substantial effects on results, as we observed on a neuron dataset collected at 5 time points from a presumably fast and synchronized process (Fig S14). This underscores the necessity for a more comprehensive understanding and modeling of cell heterogeneity, and optimal experimental design for both the number and timing of the time points to generate more informative data.

In summary, our biophysically motivated model of the dynamical processes captured in single-cell data enables the inference of process times and parameters with biophysical interpretations. It presents an alternative approach to unveil continuous latent cell representations within a well-defined and rigorous framework, and highlights the limitations of what can be inferred using current snapshot single-cell genomics data.

## 4 Methods

We begin by describing our trajectory model, followed by a description of the inference procedure. Next, we explain the analysis pipeline, including our gene and model selection procedure. Then, we elucidate the simulation setup. Finally, we detail the preprocessing of real datasets.

Throughout the Methods we denote data by **X**, and note that **X** corresponds to a cell by gene by species array. We use *i* to index cells, *j* to index genes, and *c* to index species. Parameters are denoted by *θ*, and latent variables by *z. p*(·) means a probability distribution.

### 4.1 Model

We used a simple and interpretable latent variable model for the probability distribution of the counts, with explicit biophysical meaning associated to each latent variable. We assumed cells are asynchronous, and we therefore introduced two latent variables that corresponded to the lineage and time of the cell. If other factors needed to be taken into account, more latent variables were added, for example, the stages of the cell cycle.

Therefore, in our trajectory model, the gene expression data of each cell **x** was described as a function of latent variables **z** = (*l, t*) which specify the lineage *l* and time *t* of the cell. By considering a particular parametric class of gene expression dynamics that specify the distribution *p*(**y**|**z**, *θ*) of in vivo counts **y**, and a sequencing noise model *p*(*x*|*y*), we mapped latent variable *z* into data space and arrived at an explicit formulation of data distribution that could be trained using the expectation-maximization (EM) algorithm. In other words, we assumed the counts of each cell **x** had the following density,

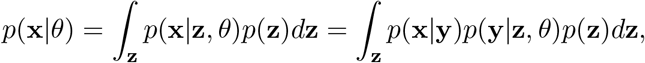

where *p*(*z*) is the distribution of latent variables.

In the following, we specify *p*(*y*|*z*, *θ*), *p*(*x*|*y*) and *p*(*z*), which are based on the transcription model, measurement model and sampling measure.

#### 4.1.1 Transcription model

Since current transcriptomic data can be used to measure the numbers of both nascent and mature transcripts, we considered the production, processing, and degradation of individual RNA molecules in our transcription model, as in previous RNA velocity literature [15].

The distribution of RNA counts in the reaction system (Eq 2) is known [22]. We assumed that the initial distribution of *U* and *S* was a product Poisson distributions, which implies that the mean parameter vector *λ* = (*λ*_*u*_, *λ*_*s*_) of *U* and *S* evolved according to Equation 3.

The functional form of *a*_*l*_(*t*) reflects our assumptions of gene expression during development. However, explicitly modeling transcription rates as functions of gene expressions is hard and tends to overfit. More fundamentally, we have some prior physical intuition; this function is actually *a*_*l*_(*λ*, **u**, *t*), where **u** is some high-dimensional vector of regulator concentrations. These data are not possible to collect using currently available technologies, although this constraint may change in the coming years. In principle, we can write down these equations, and even simulate them (using dyngen [34] or the stochastic simulation algorithm), but we strived to start with an analytically tractable model, particularly to recapitulate and formalize the increasingly popular RNA velocity framework. Therefore, we used a phenomenological model and didn’t consider gene interactions. Rather, the effect of transcriptional regulation on each gene during development was summarized into synchronized state switching and each state had its transcription rate. This formalized transient cell types.

To be able to account for multiple lineages, we assumed cell states 𝒮 = 0, 1, …, *S* formed a directed rooted tree and each lineage was a path from root 0 (initial condition) to a leaf with length *K*. The trajectory structure described the graph, and recorded the states *s*(*l, k*) ∈ 𝒮, *l* = 1, …, *L, k* = 1, …, *K* of the *L* lineages during the *K* stages. We treated the trajectory structure as known since it can typically be obtained from our previous knowledge of the data. Specifically, for lineage *l*, we assumed *a*_*l*_(*t*) is a piecewise constant function: *a*_*l*_(*t*) = *α*_*s*_(*l, k*), where the cellular state index s is determined by the lineage *l*, and the time interval *k* to which *t* belongs, i.e., *t* ∈ (*τ*_*k−*1_, *τ*_*k*_]. The state index *s* = *s*(*l, k*) was determined by the trajectory structure. For example, given the trajectory structure in Fig. 1, *s*(*l* = 1, *k* = 0) = *s*(*l* = 2, *k* = 0) = 0, *s*(*l* = 1, *k* = 1) = *s*(*l* = 2, *k* = 1) = 1, *s*(*l* = 2, *k* = 2) = 2 and *s*(*l* = 2, *k* = 2) = 3.

Then, the parametric solution of *λ* was given by

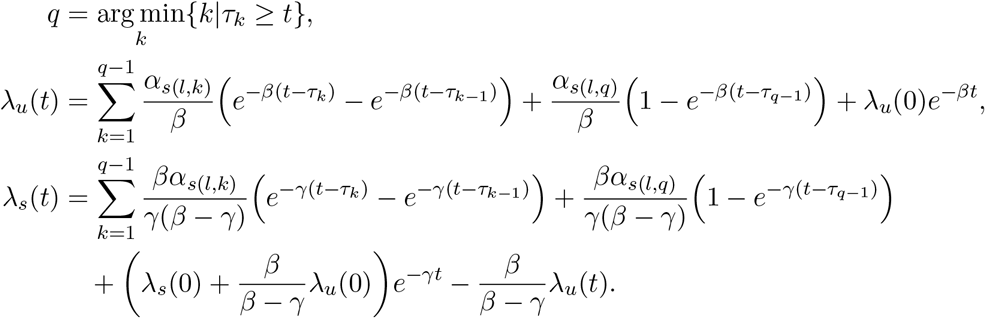

Assuming cells are at steady states at time 0, we have 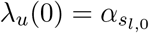 and 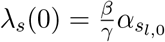. Therefore, for each gene, the parameters are *θ* = (*α*, *β, γ, τ*).

#### 4.1.2 Measurement model

Next, we needed to have a measurement model that connected counts number **y** in cells to the observed counts **x** in a single-cell RNA-seq experiment.

We assumed each molecule of mRNA produces *Bernoulli* (*q*) number of captured RNA molecules, which is also an good approximation of a Poisson model with low mean [35, 36].

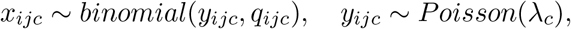

which translates to

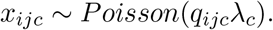

The capture rate *q*_*ijc*_ can potentially be dependent on cell read depth, genes/species-specific bias of the mRNA capture methods. In our model, we only considered the read depth (cell size) variance of cells, i.e., *q*_*ijc*_ = *c*_*i*_ where *c*_*i*_ is the read depth (cell size) of cell i. This is equivalent to multiplying the mean of Poisson distributions by a constant *c*_*i*_. Since usually only the relative value of read depth *c*_*i*_ can be available, we absorbed the mean of *c*_*i*_ into *α* and infer their product directly:

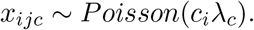

#### 4.1.3 Sampling distribution

Now that we have defined the *p*(*x*|*z*, *θ*), the only remaining thing to complete *p*(*x*|*θ*) is the sampling distribution *p*(*z*), which describes the prior distribution of latent variables. Given the formula of *p*(*y*|*z*, *θ*), it is easy to see that the model is not identifiable if *p*(*z*) is not fixed, because we can change *p*(*z*) together with *β* and *γ* easily without changing *p*(*x*), for example, by scaling *β* and *γ* and transform *p*(*z*) accordingly. Therefore, we assumed *t* ∈ [0, 1] and fixed a uniform prior for *t* on (0, 1]. With this given prior distribution, the model was identifiable, because the parameterized form of the means of Poisson distributions is identifiable and Poisson distribution is identifiable [37]. If one has information about real time, one can adjust the range and prior of *t* to be have more physical meaning. For example, for the cell cycle dataset, if one knows that the whole cycle takes 24 hours, then one can either set the range of *t* to be [0, 24], or scale the results by dividing both *β* and *γ* by 24, while keeping the other parameters unchanged.

#### 4.1.4 Connection to cluster model

In the fast dynamic limit, as there are few cells out of steady states, transitions (edge) disappear and only states (nodes) remain, and both the weight of lineages and the length of time interval (or the weight at t=0 for state 0) determine the mixture weights of clusters. Specifically, for the state 0, its weight equals the weight at t=0, while for the state 2, its weight equals the product of the length of the second time interval (*τ*_1_, *τ*_2_] and the weight of the second lineages. Thus, the Poisson mixtures model strictly belongs to the degenerate cases of our trajectory model, which connects Chronocell to biophysical cluster models like meK-Means [38], barring differences in noise models and hard/soft assignment. This does not mean that our trajectory model should be fit on cluster data, because the resulted process time is no longer meaningful. Instead, these connections allowed us to better compare the models, and determine which model was more appropriate (Section 4.3.2).

### 4.2 Inference

#### 4.2.1 Maximum likelihood estimates of parameters by EM algorithm

We used the expectation–maximization algorithm to estimate model parameters 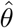. For simplicity, we discretized the latent variable *t* with finite regular grid points over the interval: *t* = *t*_1_,, *t*_*M*_, which basically approximates a continuous measure with a discrete measure. Then, the latent variable z=(l,t) describing lineage and time is discrete, and we write 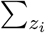 to denote the summation over all L lineages and M time grid points, i.e., 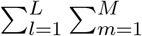. With this, the objective function becomes

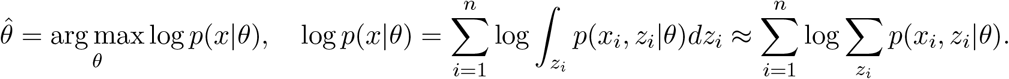

and since 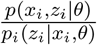 is a constant for all *z*_*i*_,

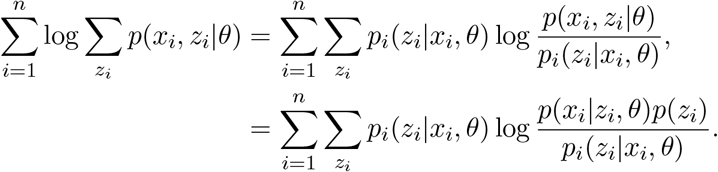

Our trajectory model makes it possible to write out *p*(*x*_*i*_|*z*_*i*_, *θ*) explicitly:

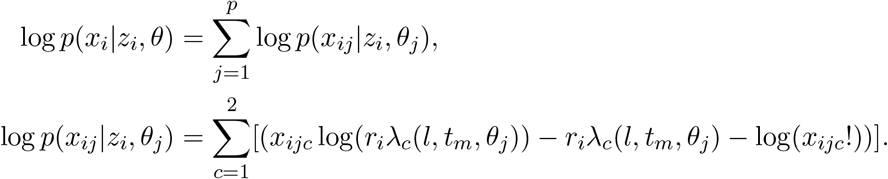

Therefore, we could use the expectation–maximization algorithm efficiently. Specifically, in the E-step of EM algorithm, we calculated the posterior distribution *p*_*i*_(*z*_*i*_|*x*_*i*_, *θ*) based on *p*(*x*_*i*_|*z*_*i*_, *θ*),

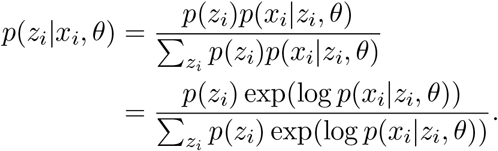

In the M-step, based on fixed *p*_*i*_(*z*_*i*_|*x*_*i*_, *θ*) and analytical form of *p*(*x*_*i*_|*z*_*i*_, *θ*), we optimized *θ*_*j*_ for each gene *j* separately by maximizing

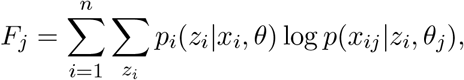

Both the value and the gradient of *F*_*j*_ can be written out analytically and computed efficiently (Sec S2). Therefore, we could use off-the-shelf quasi-Newton methods for optimizing *F*_*j*_ with respect to *θ*_*j*_, e.g., ‘L-BFGS-B’ method in minimize function provided by Scipy [39, 40].

In each step of EM algorithm, we alternated between the expectation and maximization steps. The implementation is based on defining the function that calculates *p*(*x*_*ij*_|*z*_*i*_, *θ*), so it would be easy to modify *p*(*x*_*ij*_|*z*_*i*_, *θ*) for different models in the future.

A warm start incorporating prior knowledge about data can help algorithm converge to the optimal *θ*^*\**^ quickly. If neither initial parameters nor *p*(*z*|*x*) is given, we use random initializations. Multiple runs with different starting points are used to avoid local minima. By default, we tried 100 different random initializations, and run 100 steps for each initialization both for random initialization and warm start.

#### 4.2.2 Read depth estimation

If we refer to all cell-wise variances as extrinsic noise, then it includes both sequencing noise and a part of biological sotchasticity. This extrinsic noise is shared by covariance between genes and variance of genes. Therefore, we can estimate such extrinsic noise using covariance to leave out the intrinsic noise of genes that are more biologically relevant.

Consider two species *A* and *B*. Suppose that *in vivo* their counts numbers are *Y*_*a*_ and *Y*_*b*_ with *Y*_*a*_ *∼ µ*_*a*_ and *Y*_*b*_ *∼ µ*_*b*_ where *µ*_*a*_ and *µ*_*b*_ are arbitrary distributions. Suppose additionally that during sequencing, the read depth (cell size) *c* is a random variable on [0,1], and *c* is independent of *Y*_*a*_ and *Y*_*b*_. Denote the counts we ultimately get by *X*_*a*_ and *X*_*b*_, where *X*_*a*_ and *X*_*b*_ are independent given c,*Y*_*a*_, and *Y*_*b*_, and E[*X*_*a*_|*cY*_*a*_] = *c* E[*Y*_*a*_] and E[*X*_*b*_|*cY*_*b*_] = *c* E[*Y*_*b*_].

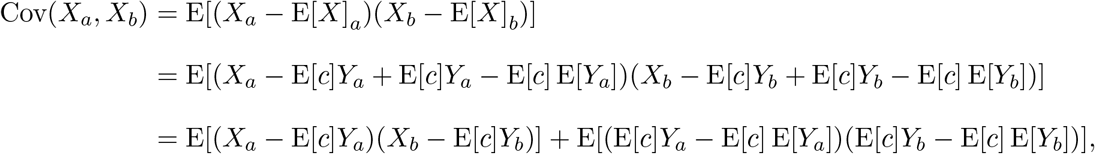

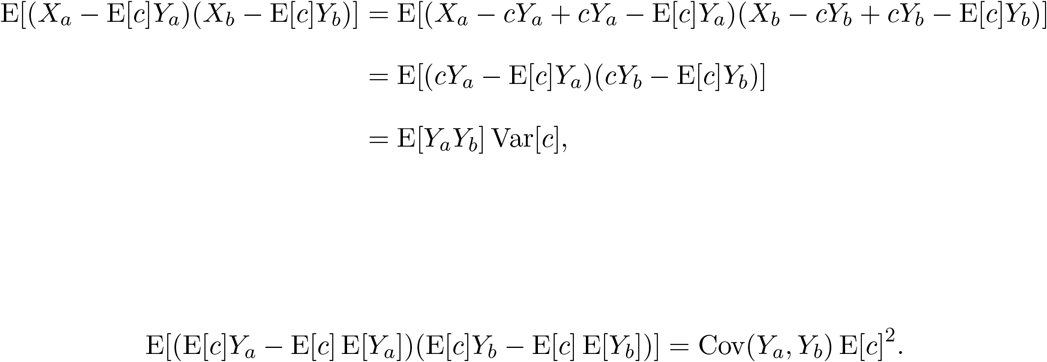

Therefore,

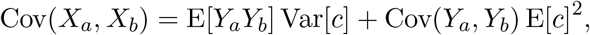

and

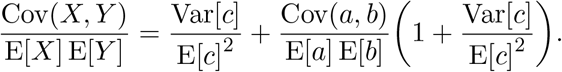

If we assume genes are independent on average, then we can estimate the CV^2^ of read depth 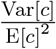 by 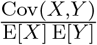. We can summarize the extrinsic noise by defining 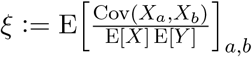, and estimate *ξ* by average normalized covariance across genes.

On the other hand, the variance of one gene is

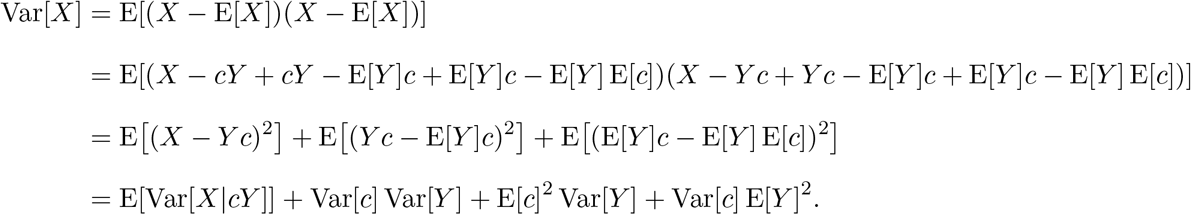

Now apply a second order approximation and suppose 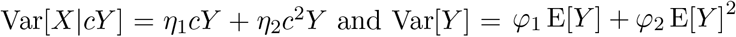 sequentially.

Then,

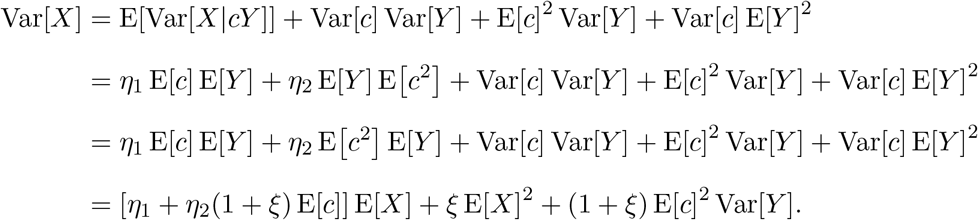

and

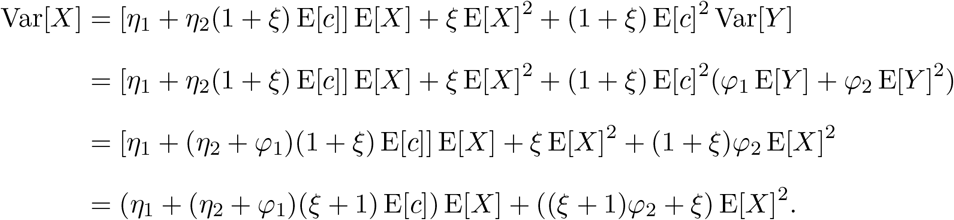

It’s easy to see that if *φ*_2_ = 0, i.e., gene intrinsic variance does not have quadratic terms, the coefficient of quadratic term of variance is also *ξ*. Therefore, if the quadratic term of the variance of a gene is closed to *ξ*, we know it is more Poissonian. In other words, it does not differentially expressed and could be used to estimate read depth.

Therefore, our procedure of read depth estimation was: first, estimate read depth CV^2^ and *ξ*; then, select genes whose quadratic term of the variance is close to *s*; finally, sum them up and normalize by the mean to calculate relative read depth. A similar, but not identical procedure of read depth estimation, was used in [41].

#### 4.2.3 Poisson mixtures model

For fitting clusters, we used Poisson mixture models. Suppose there are S mixtures with transcription rates *α*_*js*_, *s* = 1, …, *S* for each gene *j*, and each mixture is at steady state. Thus, the parameters of one gene are 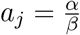 and 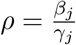, since only the ratio is identifiable.

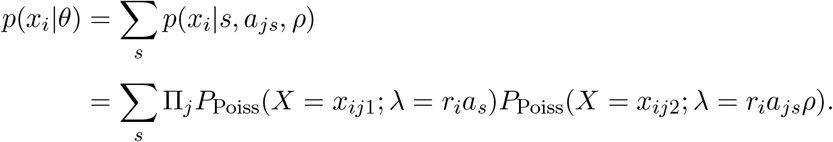

Similarly to the Gaussian mixture model, we could use the EM algorithm to infer parameters (including mixture weights) and posteriors.

### 4.3 Analysis

#### 4.3.1 Fisher information

The Fisher information matrix (FIM) is defined to be

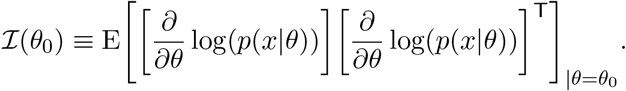

Since

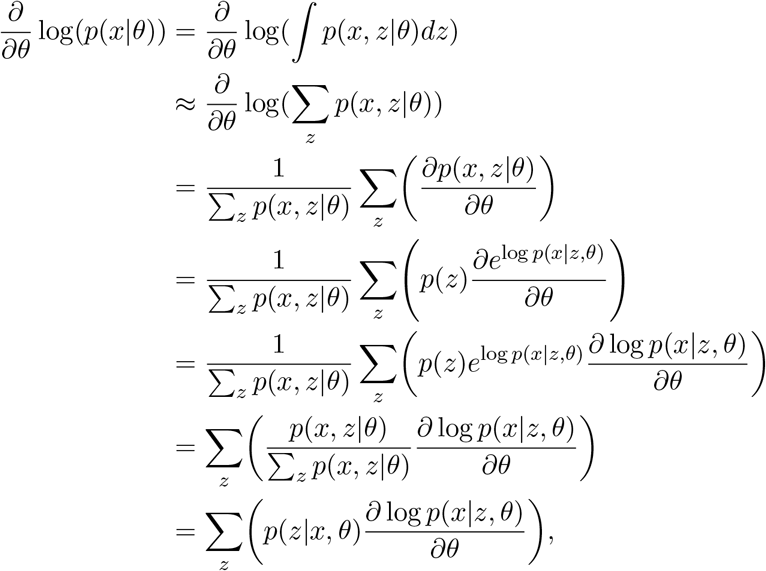

we could calculate FIM numerically with the explicit form of the derivative and the posterior distribution.

#### 4.3.2 Gene selection

We did not use absolute likelihoods to select dynamical genes, because different genes have different scales and are not directly comparable. Instead, we noticed that the cluster model serves as a natural reference point for comparison, as genes with no dynamics can be fitted equally well, if not better, by clusters. Therefore, we decided to use the relative likelihood as a criterion for gene selection.

For trajectory model,

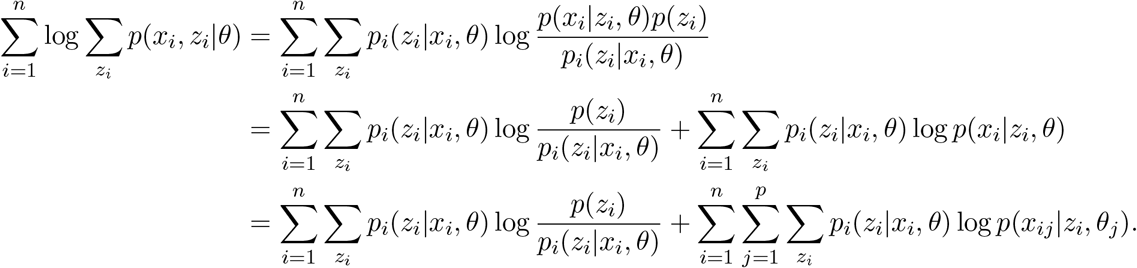

Similarly for clusters model,

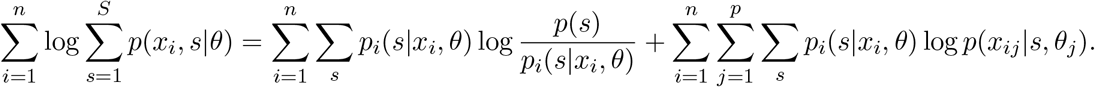

We defined the gene-wise likelihood of gene j for trajectory model to be

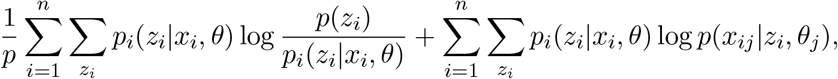

and for clusters,

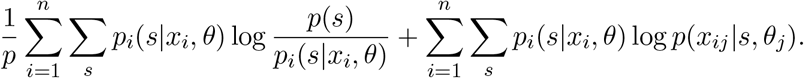

The first term is meant to represent the difference in likelihood caused by different flexibility of latent variables of two models.

For gene selection, we compared the gene likelihood of two models and selected genes whose gene likelihood of trajectory model was higher than those of the cluster model.

#### 4.3.3 Uncertainty assessment

We used uncertainty/instability to falsify results. First, we evaluated the variation of different random initializations to assess the uncertainty of the inference. This was done by performing multiple (usually 100) random initializations. Specifically, we evaluated whether the process time estimation of initializations with high ELBO scores concentrated around the correct direction. To quantify this, we classified the output of each random initialization to be correct if the process time estimates had a correlation higher than 0.8 with the reference, and we used different ELBO score thresholds to compute the precision-recall curve, which were then summarized by average precision (AP). An ideal case with high ELBO scores concentrated around correlation one led to an AP close to one while multiple comparable maxima lead to low AP.

Another approach to assess uncertainty is through bootstrap resampling. The same inference procedure is applied to the resampled data, and the variation in the resultant process time serves as an indicator of instability. Specifically, we calculate the correlation of process time between the original and the resampled data and interpret instability as large variance of the correlation.

### 4.4 Simulations

We randomly generated parameters for 200 genes and sampled 2000 cells by default unless otherwise specified. Cells were sampled uniformly over process time and lineages. We then fitted the model with the correct trajectory structure and synchronized model if not stated otherwise.

For the simulation parameters, we assumed that the transcription parameters (*α*,*β,γ*) followed the log-normal distribution *lognormal*(*µ*, σ), where *µ* is the mean and σ is the standard deviation of the variable’s natural logarithm. We attempted to derive realistic parameters for the distributions from the literature. Hence, we assumed *β ∼ lognormal*(2, 0.5), *γ ∼ lognormal*(0.5, 0.5) so that unspliced to spliced ratio was around 0.2 and their distributions resembled those determined from metabolic labeling datasets [24]. The *α* were assumed to follow *lognormal*(2, 1), so that the mean of spliced counts were similar to those in scRNA-seq data. We assumed the read depth follows Beta distribution 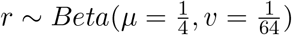, where *µ* was the mean and *v* the variance. For simulations with Gamma noise, we multiplied the mean of Poisson distribution *λ* by a random variable following Gamma distribution with mean 1 and variance 0.5.

For simulations under the desynchronized model, gene-wise *τ*_*k*_ was sampled from a uniform distribution on 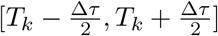, where *T*_*k*_ corresponds to the global *τ*_*k*_ in the synchronized model, and *Δτ* was the smallest interval length ensuring that *τ*_*k*_ retains its order.

To vary the sampling distributions, we generated a Gaussian distribution with a random mean and standard deviation 0.05. We then sampled both from the Gaussian and the uniform distribution, and blended them together in different proportions. The percentages of time sampled from a Gaussian ranged from 0 to 1, increasing by 0.1 increments.

### 4.5 Real datasets preprocessing

To estimate the squared coefficient of variation of the read depth 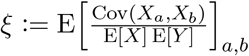, we calculated the covariance matrix of all genes with nonzero means, which was divided by the mean squared and averaged across gene pairs to calculate the mean normalized covariance as an estimate of *ξ*. We then selected Poissonian genes whose variances are close to baseline variance with reasonably large mean (*var* < 1.2(*µ* + *ξµ*^2^), *µ* > 0.01). However, as some genes can be co-regulated and, therefore, correlated, we calculated the mean normalized covariance of Poissonian genes and repeated selected new Poissonian genes until the mean normalized covariance no longer changed. This typically occurred after two iterations. Finally, we normalized the sum of counts of the selected Poissonian genes by their mean to obtain the relative read depth estimates, which were then used as fixed parameters during the fitting process.

Based on simulations of factors of informativeness of the data, including numbers of cells and genes, the average mean of *α* parameters as well as *β* and *γ* ratio, we determined the procedure of filtering genes for fitting: we applied count mean thresholds (0.02 for unspliced and 0.1 for spliced), filtered out genes with a small unspliced to spliced ratio 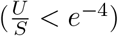, and finally, selected genes with a variance larger than 1.2 times the baseline variance, i.e., *var* > 1.2(*µ* + *ξµ*^2^), where *µ* is the mean, *var* is the variance, and *ξ* is the read depth CV^2^. For cell cycle data, we also constrained genes to the Gene Ontology term “cell_cycle” (GO:0007049). We occasionally adjusted the variance threshold to 1.5 in order to end up with 50–200 genes. Then trajectory and Poisson mixture models were fitted on those genes.

## 5 Data and Code Availability

The data sets analyzed in this paper are from previously published research, and are publicly available. The 10x Genomics 10k Human PBMC data can be downloaded from https://zenodo.org/record/7388133/files/GP_2021_3_raw_data.tar.gz?download=1 with metadata from https://doi.org/10.22002/7fbwg-rx843. The forebrain dat can be downloaded from http://pklab.med.harvard.edu/velocyto/hgForebrainGlut/hgForebrainGlut.loom. The erythroid data can be downloaded from https://ndownloader.figshare.com/files/27686871. The RPE1 and Neuron data can be downloaded from the dynamo packages https://github.com/aristoteleo/dynamo-release, or directly from https://www.dropbox.com/s/25enev458c8egn7/rpe1.h5ad?dl=1 and https://www.dropbox.com/s/g1afqdcsczgyj2m/neuron_splicing_4_11.h5ad?dl=1 respectively.

All code used to generate the results and figures in the paper is available at https://github.com/pachterlab/FGP_2024.

## Declarations

GG is currently an employee of Fauna Bio. This work was completed while GG was at the California Institute of Technology.

## 6 Acknowledgements

We thank Maria Carilli, Tara Chari, and Catherine Felce for helpful discussions and feedback on the manuscript.

