## Supplementary notes for "Trajectory inference from single-cell genomics data with a process time model"

Meichen Fang[1], Gennady Gorin[2], and Lior Pachter[1,3,*]

[1]Division of Biology and Biological Engineering, California Institute of Technology, Pasadena, CA, USA

[2]Fauna Bio, Emeryville, CA, USA

[3]Department of Computing and Mathematical Sciences, California Institute of Technology, Pasadena, CA, USA

#### S1 Challenges with the pseudotime concept

##### S1.1 Trajectory methods overview

Single-cell genomics trajectory inference methods have mostly relied on similarity metrics: distance based methods reconstruct the trajectory based on some distance metrics in gene expression space under the assumption that cells that are more similar in gene expression space are also closer in pseudotime [1, 2, 3]. Manifold-learning based methods draw the trajectories in a reduced dimension space based on connectivity, i.e., similarity [4, 5]. Probability/Markov chain based methods also calculate transition probabilities based on distances [6]. However, pseudotime based on similarity/distance is inherently descriptive and unable to be extended to reflect physical meaning, because state spaces of dynamical processes are not isotropic. There are a few exceptions that implicitly define generative models of single-cell RNA-seq (scRNA-seq) data with pseudotime, modeling dynamics of gene expression along differentiation processes in a way that can be reformulated as driven by cell states switching models [7]. These ideas have motivated our model.

One important observation is that the usage of Markov Chain model in trajectory inference, as well as other single cell analysis, can be fundamentally flawed. This is because frequently cells are samples drawn from different chains, instead of a sequence of observations of one single chain. The Markov chain formalism is therefore not applicable to single cell studies without making further assumptions.

#### S1.2 Circularity in pseudotime-based analysis

Due to the exploratory nature of trajectory inference, all variables are used to fit the model, but not all variables are informative. Specifically, it is natural to assume sparsity and focus on a few "marker genes" in downstream analysis. At present, a multi-stage method is commonly employed for pseudotime-based analysis. In the initial stage, trajectory and pseudotime are fitted, followed by the second stage, where hypothesis testing is utilized to select genes that are variable along trajectories.

Ideally, with a predefined and well-parameterized model, we can construct confidence intervals for parameters using Bayesian methods or the bootstrap. However, interpretation can be difficult even for PCA loadings [8]. As heuristic methods become more popular, the variable selection problem is highly entangled with model construction, and the question of how to perform valid inference is not straightforward. Current methods usually first perform trajectory inference, and then test whether genes expression have dependency with pseudotime using the same dataset. It is well-known that such tests are not valid and can lead to inflated false positive rates [4, 9, 10, 11]. The same issue for clustering has also been well discussed in several recent studies [12, 13, 14]. Here we briefly summarize the issue in the context of trajectory inference.

First, fitting and testing using the same dataset means that one has cherry-picked the most significant association and results are consequently biased upward, which is known as post-selection inference [15, 16]. Furthermore, even if one uses separate datasets for fitting and testing, there is an inherent circularity in the hypothesis testing. Specifically, during trajectory inference, one selects a transformation (that defines a trajectory) $\hat{f}$ that maps a cell to a pseudotime based on its gene expression. Then, by testing whether genes expression associates with pseudotime, one is asking whether $x$ associates with $\hat{f}(x)$, which just echoes the model fitted and does not perform the hypothesis testing validly.

#### S1.3 PCA as an example

To see this circularity, consider a single component model where trajectory is replace by first component of PCA. Denote the data by $X$, and let Y be the normalized data matrix, i.e. $Y_{ij} = X_{ij} - \bar{X}_j$ for covariance PCA and $Y_{ij} = \frac{X_{ij} - \bar{X}_j}{\sqrt{\sum_i (X_{ij} - \bar{X}_j)^2}}$ for correlation PCA. Write $S = \frac{1}{n} Y^T Y = V \Lambda V^T$ and denote the first eigenvector in PCA by $v$ (first column of V) and first eigenvalue by $\lambda$. Then for first principal component scores, which are the latent variable $z$ we want, we have $z = Yv$ and $\frac{1}{n} y_j^T z = \frac{1}{n} y_j^T Y v = \frac{1}{n}(Y^T Y v)_j = \lambda v_j$. If we directly perform linear regression of $z$ on the expression of mean-centered gene $y_j$, the slope is $v_j$.

Even after data splitting, if we follow the normal linear regression procedure and test the null hypothesis that $\beta = 0$, we will derive a t-statistic that is still biased. The correct way is to account for the projection $z = Yv$ and test $\beta = v_j$.

#### S1.4 Current solutions

In practice, there are only a few papers that have taken this circularity into account. One possible solution is count splitting if counts number are high enough [17]. Another possible solution is data splitting, where we split the dataset into two parts, select our model using the first part and do the inference using the second. Specifically, to perform rigorous hypothesis testing and get some valid p-value, we can perform permutation test. However, it means that we need to generate sets of permuted data and perform the whole procedure of trajectory inference and DE analysis on each set. This approach does not seem to have been explored or adopted in any currently used tools.

More crucially, this kind of pseudotime-based analysis only answers data analytic questions, i.e., data summary and analysis. They are typically not concerned with goodness of fit/model selection, thus providing no information about the correctness of the fitted model.

#### S2 Analytical solution of Poisson mean and its derivative

Assume cells starting from a steady state, then $\lambda_u(0) = \frac{\alpha_0}{\beta} := a_0$ and $\lambda_s(0) = \frac{\alpha_0}{\gamma}$. Denote $\mathbb{1}_k(t) = [t \geq \tau_k]$. Let $\alpha(t) = \alpha_{s(l,q)}$ where $q = \min_k\{k|\tau_k \geq t\}$, and let $a_s = \frac{\alpha_s}{\beta}$.

Then for lineage l, time $t_m$ and gene j with parameters $\theta_j = (\alpha, \beta, \gamma, \tau)$,

$$\begin{aligned}
\lambda_u(l, t_m, \theta_j) &= \sum_{k=1}^{K} a_{s(l,k)} \left( e^{-\beta_j \mathbb{1}_k(t)(t_m - \tau_k)} - e^{-\beta_j \mathbb{1}_{k-1}(t)(t_m - \tau_{k-1})} \right) + a_0 e^{-\beta_j \mathbb{1}_0(t)(t_m - \tau_0)}, \\
\lambda_s(l, t_m, \theta_j) &= \frac{\beta^2}{\gamma(\beta - \gamma)} \left[ \sum_{k=1}^{K} a_{s(l,k)} \left( e^{-\gamma \mathbb{1}_k(t)(t_m - \tau_k)} - e^{-\gamma \mathbb{1}_{k-1}(t - \tau_{k-1})} \right) + a_0 e^{-\gamma_j \mathbb{1}_0(t)(t_m - \tau_0)} \right] \\
&\quad - \frac{\beta}{\beta - \gamma} \lambda_u(l, t_m, \theta_j).
\end{aligned}$$

Since

$$\frac{\partial F}{\partial \theta} = \sum_{i=1}^{n} \sum_{l=1}^{L} \sum_{m=1}^{M} q_i(l, t_m) \sum_{c=u,s} \left( \frac{x_{ijr}}{\lambda_c(l, t_m, \theta_j)} - 1 \right) \frac{\partial \lambda_c(l, t_m, \theta_j)}{\partial \theta},$$

in order to calculate the derivative of $F$ with respect to $\theta$, we need to calculate the derivatives of $y$ with respect to $\theta$.

$$
\begin{aligned}
\frac{\partial \lambda_u}{\partial a_0} &= e^{-\beta_j \mathbb{1}_0(t)(t_m - \tau_0)}, \\
\frac{\partial \lambda_u}{\partial a_{s(l,k)}} &= e^{-\beta_j (t_m - \tau_k)\mathbb{1}_k(t)} - e^{-\beta_j (t_m - \tau_{k-1})\mathbb{1}_{k-1}(t)}, \\
\frac{\partial \lambda_u}{\partial \beta} &= \sum_{k=1}^{K} a_{s(l,k)} \Big( -(t_m - \tau_k)\mathbb{1}_k(t) e^{-\beta_j (t_m - \tau_k)\mathbb{1}_k(t)} + \mathbb{1}_{k-1}(t)(t_m - \tau_{k-1}) e^{-\beta_j (t_m - \tau_{k-1})\mathbb{1}_{k-1}(t)} \Big) \\
&\quad - a_0 \mathbb{1}_0(t)(t_m - \tau_0) e^{-\beta_j \mathbb{1}_0(t)(t_m - \tau_0)}, \\
\frac{\partial \lambda_s}{\partial a_{s(l,k)}} &= \frac{\beta^2}{\gamma(\beta - \gamma)} \frac{\partial \lambda_u}{\partial a_{s(l,k)}} - \frac{\beta}{\beta - \gamma} \frac{\partial \lambda_u}{\partial a_{s(l,k)}}, \\
\frac{\partial \lambda_s}{\partial \beta} &= \frac{\beta^2 - 2\beta\gamma}{\gamma(\beta - \gamma)^2} \left[ \sum_{k=1}^{K} a_{s(l,k)} \Big( e^{-\gamma \mathbb{1}_k(t)(t_m - \tau_k)} - e^{-\gamma \mathbb{1}_{k-1}(t - \tau_{k-1})} \Big) + a_0 e^{-\gamma \mathbb{1}_0(t)(t_m - \tau_0)} \right] \\
&\quad - \frac{\beta}{\beta - \gamma} \frac{\partial \lambda_u}{\partial \beta} + \frac{\gamma}{(\beta - \gamma)^2} \lambda_u, \\
\frac{\partial \lambda_s}{\partial \gamma} &= \frac{-\beta^3 + 2\beta^2\gamma}{\gamma^2(\beta - \gamma)^2} \left[ \sum_{k=1}^{K} a_{s(l,k)} \Big( e^{-\gamma \mathbb{1}_k(t)(t_m - \tau_k)} - e^{-\gamma \mathbb{1}_{k-1}(t - \tau_{k-1})} \Big) + a_0 e^{-\gamma \mathbb{1}_0(t)(t_m - \tau_0)} \right] \\
&\quad + \frac{\beta^2}{\gamma(\beta - \gamma)} \sum_{k=1}^{K} a_{s(l,k)} \Big( -\mathbb{1}_k(t)(t_m - \tau_k) e^{-\gamma \mathbb{1}_k(t)(t_m - \tau_k)} + \mathbb{1}_{k-1}(t)(t - \tau_{k-1}) e^{-\gamma \mathbb{1}_{k-1}(t - \tau_{k-1})} \Big) \\
&\quad - \frac{\beta^2}{\gamma(\beta - \gamma)} \mathbb{1}_0(t)(t_m - \tau_0) a_0 e^{-\gamma \mathbb{1}_0(t)(t_m - \tau_0)} - \frac{\beta}{(\beta - \gamma)^2} \lambda_u.
\end{aligned}
$$

$\tau_k$ can be different for different genes in desynchronized model, and their derivatives are:

$$
\begin{aligned}
\frac{\partial \lambda_u}{\partial \tau_k} &= \big( a_{s(l,k)} - a_{s(l,k+1)} \big) \mathbb{1}_k(t) \beta e^{-\beta \mathbb{1}_k(t)(t - \tau_k)}, \\
\frac{\partial \lambda_s}{\partial \tau_k} &= \big( a_{s(l,k)} - a_{s(l,k+1)} \big) \mathbb{1}_k(t) \frac{\beta^2}{(\beta - \gamma)} e^{-\gamma \mathbb{1}_k(t)(t - \tau_k)} - \frac{\beta}{\beta - \gamma} \frac{\partial \lambda_u}{\partial \tau_k}.
\end{aligned}
$$

### References


[1] Laleh Haghverdi, Maren Büttner, F Alexander Wolf, Florian Buettner, and Fabian J Theis. Diffusion pseudotime robustly reconstructs lineage branching. *Nat. Methods*, 13(10):845–848,

October 2016.

[9] David Lähnemann, Johannes Köster, Ewa Szczurek, Davis J McCarthy, Stephanie C Hicks, Mark D Robinson, Catalina A Vallejos, Kieran R Campbell, Niko Beerenwinkel, Ahmed

Mahfouz, Luca Pinello, Pavel Skums, Alexandros Stamatakis, Camille Stephan-Otto Attolini, Samuel Aparicio, Jasmijn Baaijens, Marleen Balvert, Buys de Barbanson, Antonio Cappuccio, Giacomo Corleone, Bas E Dutilh, Maria Florescu, Victor Guryev, Rens Holmer, Katharina Jahn, Thamar Jessurun Lobo, Emma M Keizer, Indu Khatri, Szymon M Kielbasa, Jan O Korbel, Alexey M Kozlov, Tzu-Hao Kuo, Boudewijn P F Lelieveldt, Ion I Mandoiu, John C Marioni, Tobias Marschall, Felix Mölder, Amir Niknejad, Lukasz Raczkowski, Marcel Reinders, Jeroen de Ridder, Antoine-Emmanuel Saliba, Antonios Somarakis, Oliver Stegle, Fabian J Theis, Huan Yang, Alex Zelikovsky, Alice C McHardy, Benjamin J Raphael, Sohrab P Shah, and Alexander Schönhuth. Eleven grand challenges in single-cell data science. *Genome Biol.*, 21(1):31, February 2020.

[10] Zhicheng Ji and Hongkai Ji. TSCAN: Pseudo-time reconstruction and evaluation in single-cell RNA-seq analysis. *Nucleic Acids Res.*, 44(13):e117, July 2016.

[11] Sophie Tritschler, Maren Büttner, David S Fischer, Marius Lange, Volker Bergen, Heiko Lickert, and Fabian J Theis. Concepts and limitations for learning developmental trajectories from single cell genomics. *Development*, 146(12), June 2019.

[12] Jesse M Zhang, Govinda M Kamath, and David N Tse. Valid post-clustering differential analysis for Single-Cell RNA-Seq. *Cell Syst*, 9(4):383–392.e6, October 2019.

[13] Lucy L Gao, Jacob Bien, and Daniela Witten. Selective inference for hierarchical clustering. December 2020.

[14] Yiqun T Chen and Daniela M Witten. Selective inference for k-means clustering. March 2022.

[15] Jonathan Taylor and Robert J Tibshirani. Statistical learning and selective inference. *Proc. Natl. Acad. Sci. U. S. A.*, 112(25):7629–7634, June 2015.

[16] Arun K Kuchibhotla, John E Kolassa, and Todd A Kuffner. Post-selection inference. *Annu. Rev. Stat. Appl.*, March 2022.

[17] Anna Neufeld, Lucy L Gao, Joshua Popp, Alexis Battle, and Daniela Witten. Inference after latent variable estimation for single-cell RNA sequencing data. *Biostatistics*, December 2022.
