## Supplementary figures for "Trajectory inference from single-cell genomics data with a process time model"

Meichen Fang[1], Gennady Gorin[2], and Lior Pachter[1,3,*]

[1]Division of Biology and Biological Engineering, California Institute of Technology, Pasadena, CA, USA

[2]Fauna Bio, Emeryville, CA, USA

[3]Department of Computing and Mathematical Sciences, California Institute of Technology, Pasadena, CA, USA

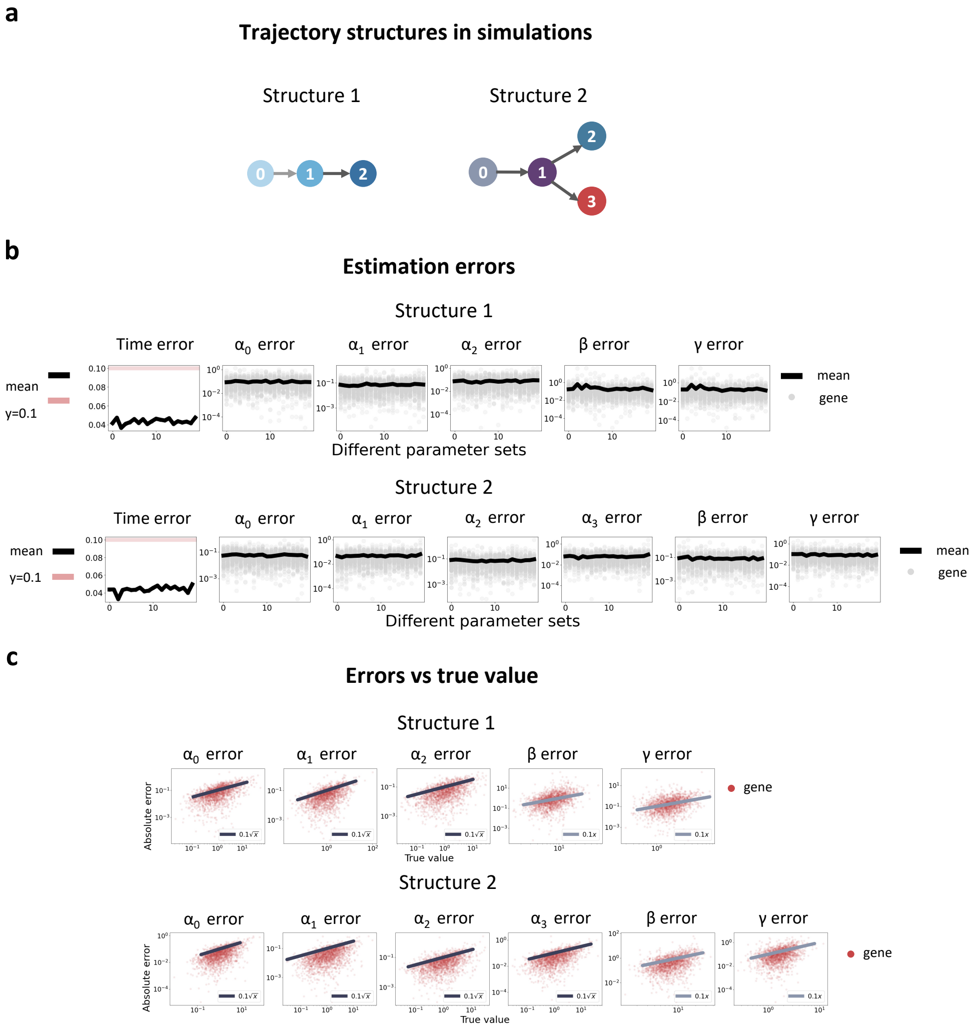


Figure S1: **Inference accuracy tested on simulation**. **a**) The two trajectory structures used in simulations. **b**) Estimation errors of different parameter sets. For time, error is root mean square error. For $\alpha, \beta, \gamma$, error is mean normalized error as described in the text. **c**) Absolute errors with respect to the true values of parameters.

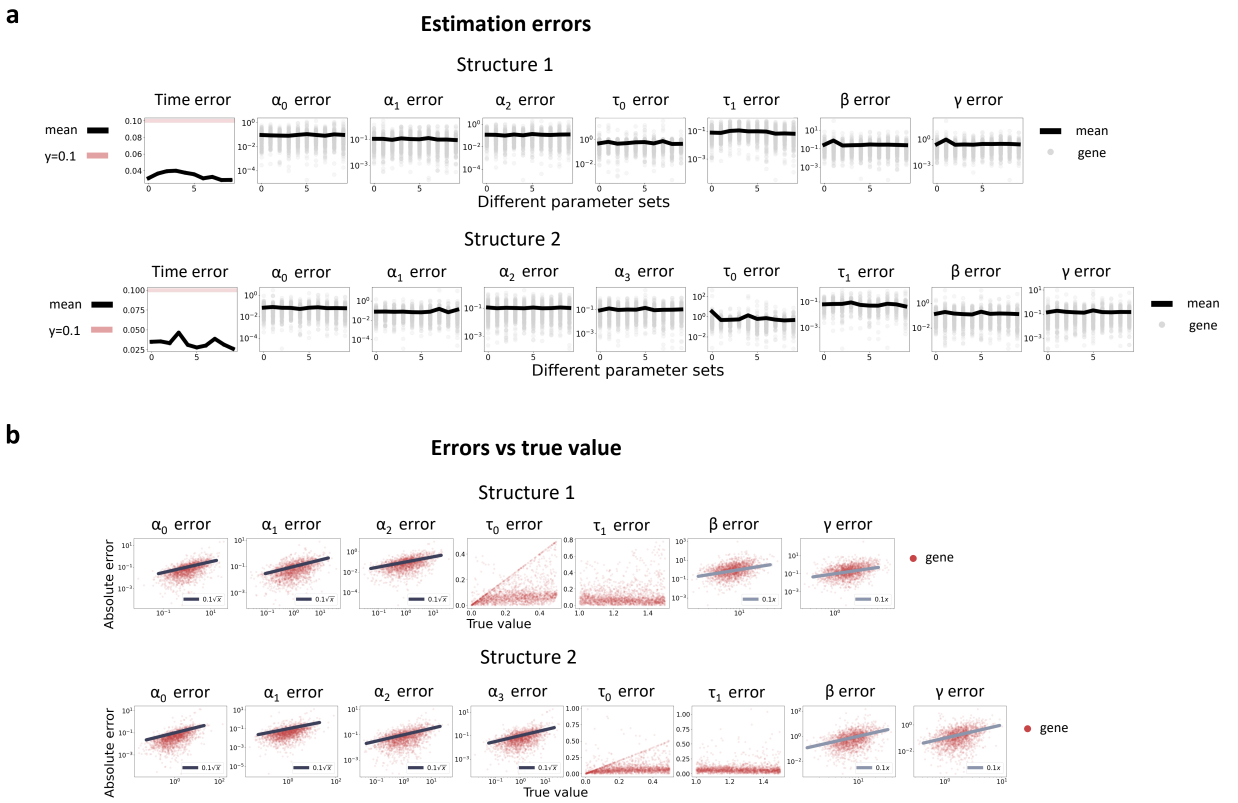


Figure S2: **Inference accuracy of desynchronized model**. Estimation errors of the desynchronized model tested on simulations with trajectory structures in Fig S1a. **a**) Errors of different parameter sets. For time, error is root mean square error. For $\alpha, \beta, \gamma$, error is mean normalized error as described in the text. **b**) Absolute errors with respect to the true values of parameters.

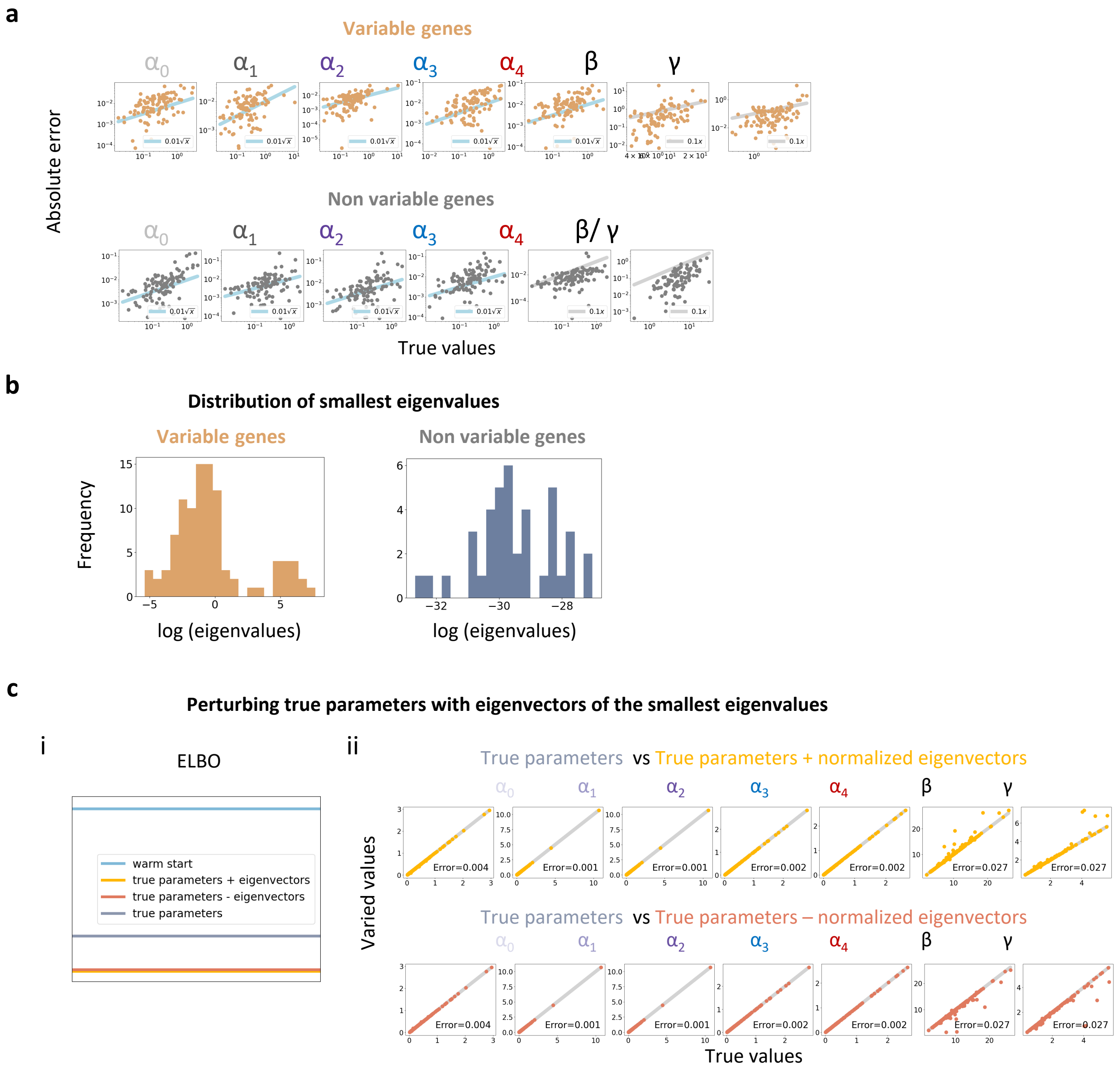


Figure S3: **Supplementary figures for demonstration of inference on simulation**. **a**) Absolute errors with respect to the true values of parameters. **b**) Distribution of the smallest eigenvalues of the Fisher information matrix of each gene. **c**) Marginal likelihood (ELBO) and varied parameters compared to true parameters. The difference between varied parameters and true parameters are the eigenvectors corresponding to the smallest eigenvalues of the Fisher information matrix, divided by the square root of the respective eigenvalues, specific to variable genes.

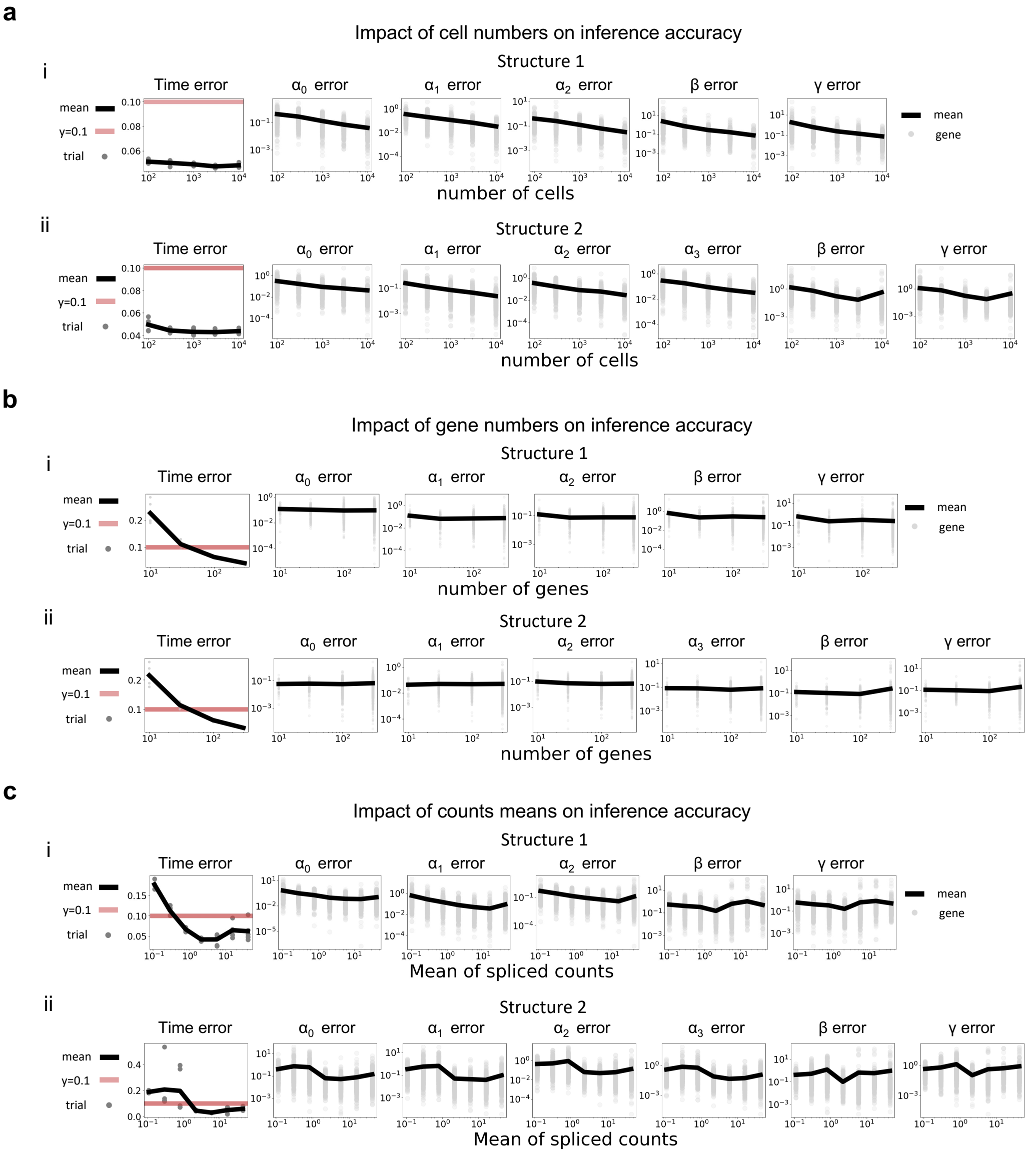


Figure S4: **Impact of cell, gene numbers, and counts means on Inference accuracy**. The trajectory structures are the same as in Fig S1a. For $\alpha, \beta, \gamma$, error is mean normalized error as described in the text. **a**) Estimation errors of different cell numbers. **b**) Estimation errors of different gene numbers. **c**) Estimation errors of different counts means due to different $\alpha$ means. For time, error is root mean square error.

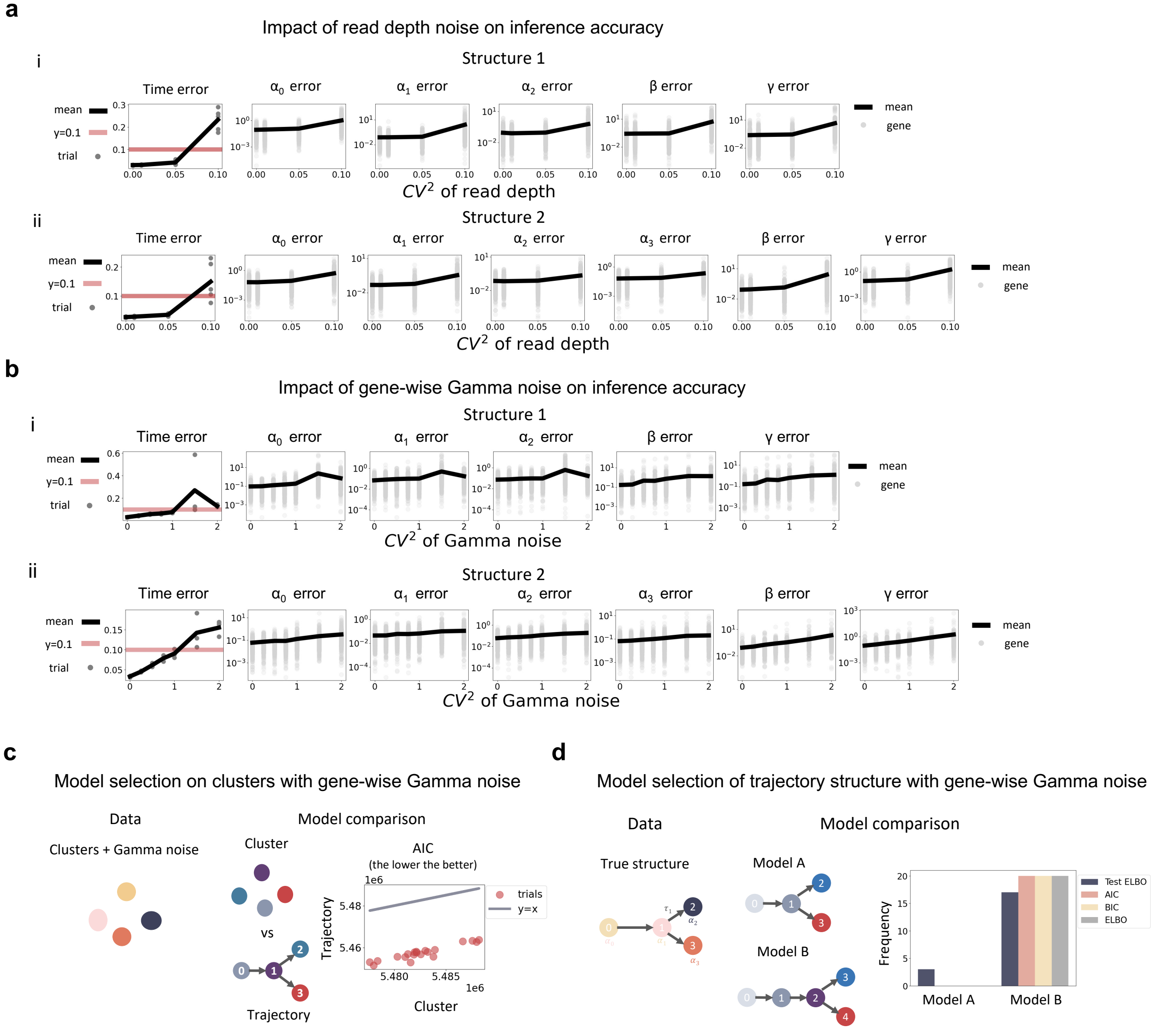


Figure S5: **Impact of noise on inference accuracy and model selection**. The trajectory structures are the same as in Fig S1a. For time, error is root mean square error. For $\alpha, \beta, \gamma$, error is mean normalized error as described in the text. **a**) Estimation errors as read depth noise increases. **b**) Estimation errors as gene-wise Gamma noise increases. **c**) Impact of gene-wise Gamma noise on model selection on clusters data. Same as in Fig. 2 except Gamma noise with $\mathrm{CV}^2$ 1 was added to Poisson mixtures to generate simulation data (Sec 4.4). **c** Impact of gene-wise Gamma noise on model selection of trajectory structure. Same as in Fig. 3j except Gamma noise with CV 1 was added in simulation.

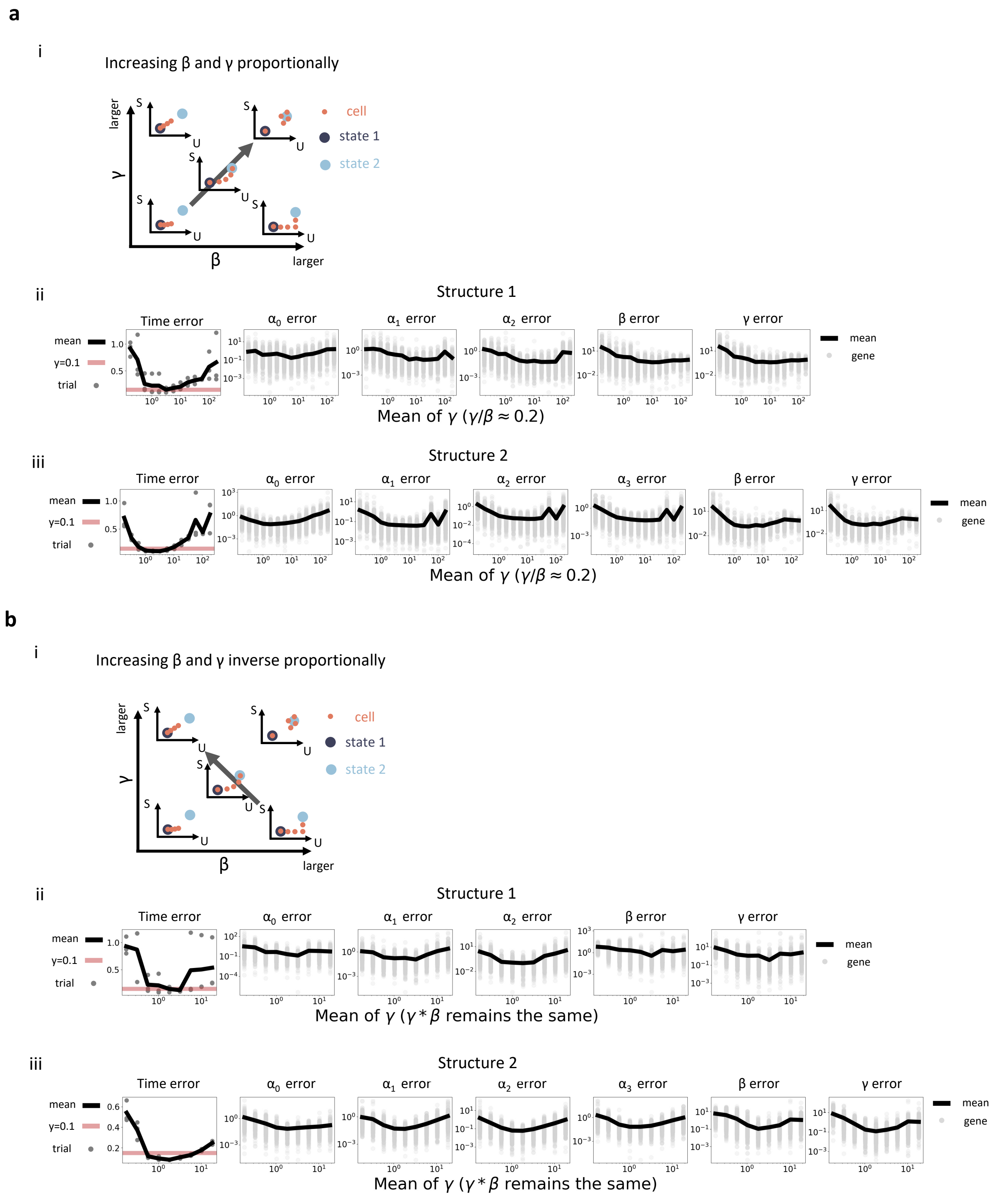


Figure S6: **Impact of dynamic timescale on inference accuracy**. The trajectory structures are the same as in Fig S1a. For time, error is root mean square error. For $\alpha, \beta, \gamma$, error is mean normalized error as described in the text. **a**) **i** Schematics of phase plots with increasing timescale. **ii** Estimation errors as time scale increases. **b**) **i** Schematics of phase plots with the ratio of $\gamma$ to $\beta$ increasing while keeping their product constant. **ii** Estimation errors as $\frac{\gamma}{\beta}$ increases.

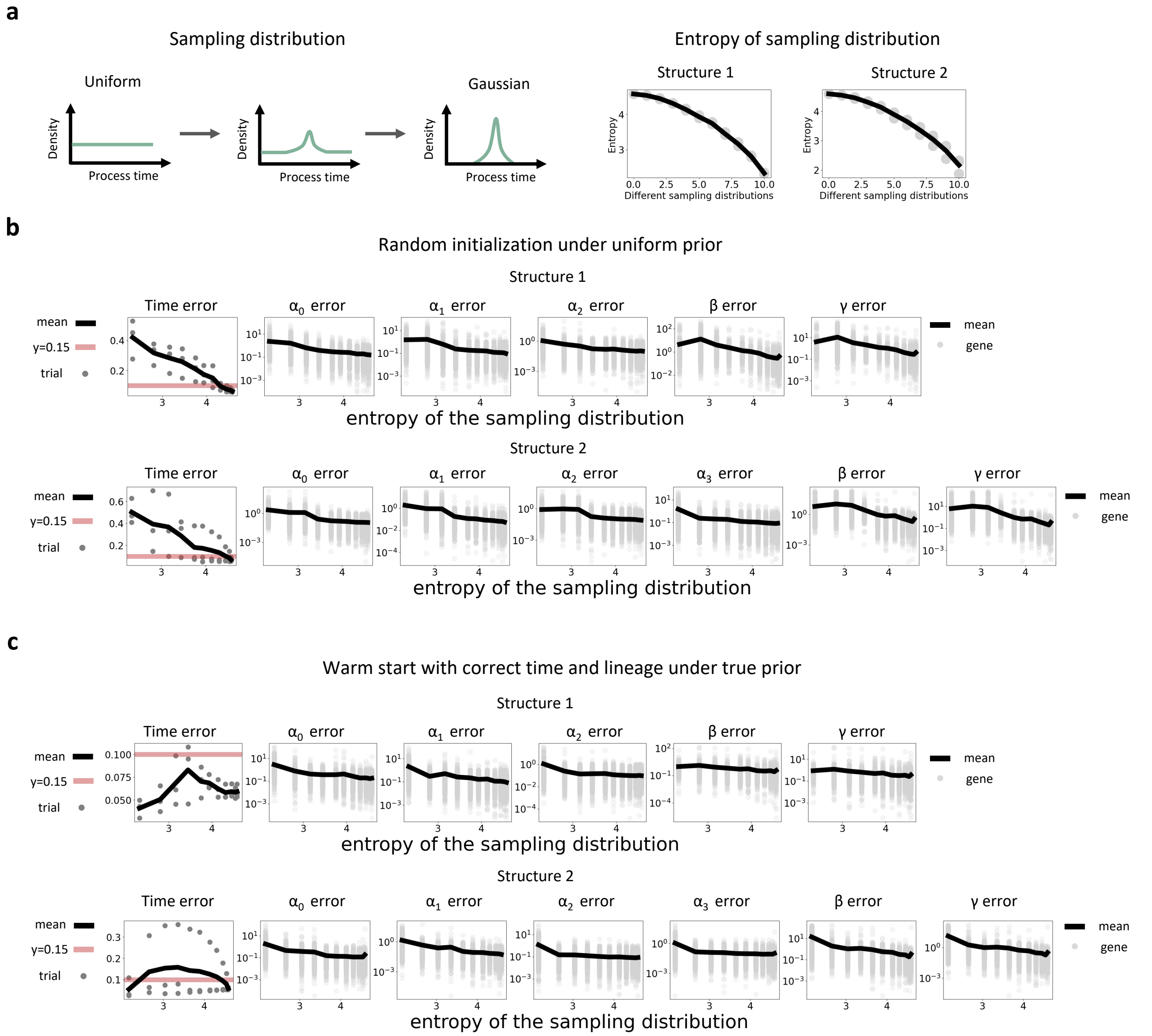


Figure S7: **Impact of sampling distribution uniformity on inference accuracy**. The trajectory structures are the same as in Fig. S1a. For time, error is root mean square error. For $\alpha, \beta, \gamma$, error is mean normalized error as described in the text. **a**) Schematics of sampling distributions used in simulation with decreasing uniformity. The sampling distributions were gradually changing from uniform distribution to Gaussian distribution. Right plot shows the entropy of the sampling distributions. **b**) Estimation errors as uniformity decreases under uniform prior. **c**) Estimation errors as uniformity decreases warm started with correct position under true prior. Fitting was initialized with posteriors calculated under true parameters, and empirical distribution of process time of samples were provided as prior for the sampling distribution.

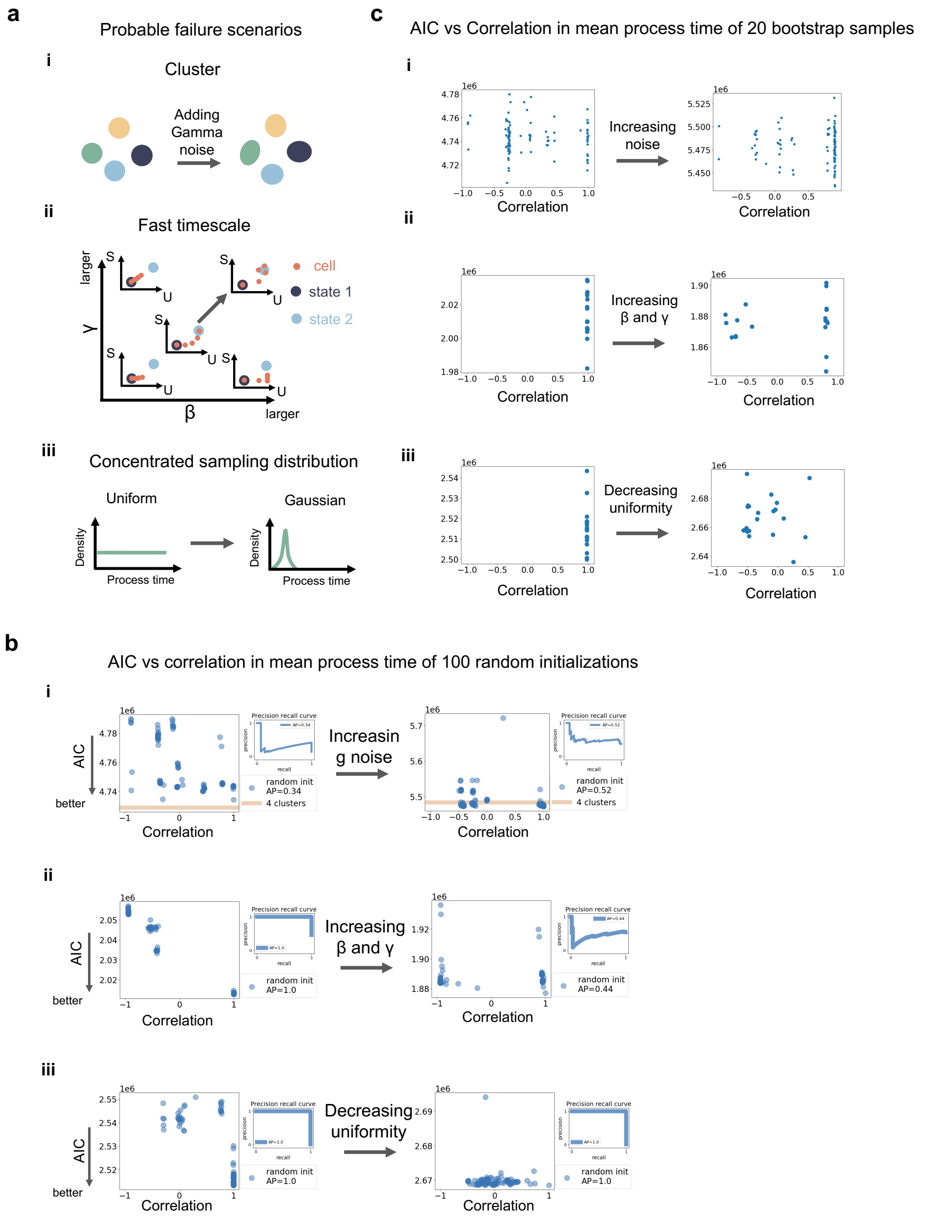


Figure S8: **Uncertainty and lack of robustness as an indicator of failure scenarios**. **a**) Schematics of three probable failure factors: clusters data, fast timescale, concentrated sampling distribution. Two example simulations were used for each case and the gray arrows indicate the their difference. **i** The two simulations are the cluster data in Fig 2 and noisy cluster data in Fig S5c respectively. **ii** The two simulations are the 5th and 13th instances of structure 1 in Fig S6b. **iii** The two simulations are the 1st and 11th instances of structure 2 in Fig S7b.**b)** AIC vs. correlation of mean process time of 100 random initializations in different scenarios. Results of two example simulations were showed. The x axis is the correlation of mean process time between each initialization and the best one. The gray arrows correspond to those in **a**). **c**) AIC vs correlation of mean process time of 20 bootstrap samples in different scenarios. Results of the two example simulations were presented in the same position as in **b**. The x axis is the correlation of mean process time between each bootstrap sample and the original one (which is the best one in **b**). The gray arrows correspond to those in **a**).

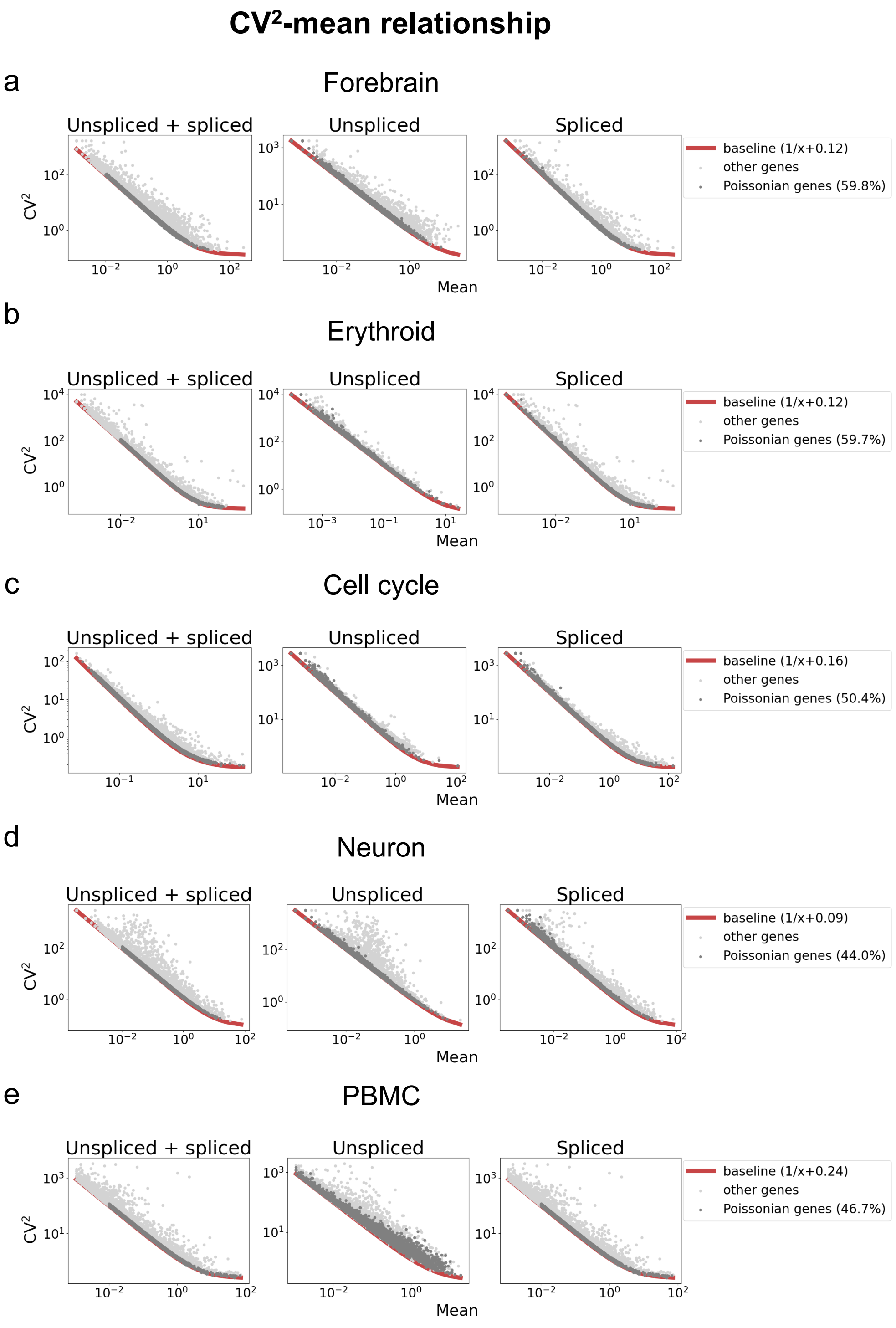


Figure S9: **Read depth estimation using normalized covariance for highly variable gene selection**. **a**) $CV^2$-mean relationship of total, unspliced and spliced counts of all genes. The baseline (red line) of variance is $1/\text{mean}$ + normalized covariance. Genes whose variance is within 1.2 times baseline are considered to be Poissonion genes (dark gray). Others are colored in light gray. **b**) Read depth estimation based on total counts of Poissonion genes. Cells are colored by their cell types. **c** Selected genes (black) for fitting plotted in the same $CV^2$-mean plot as in **a**).

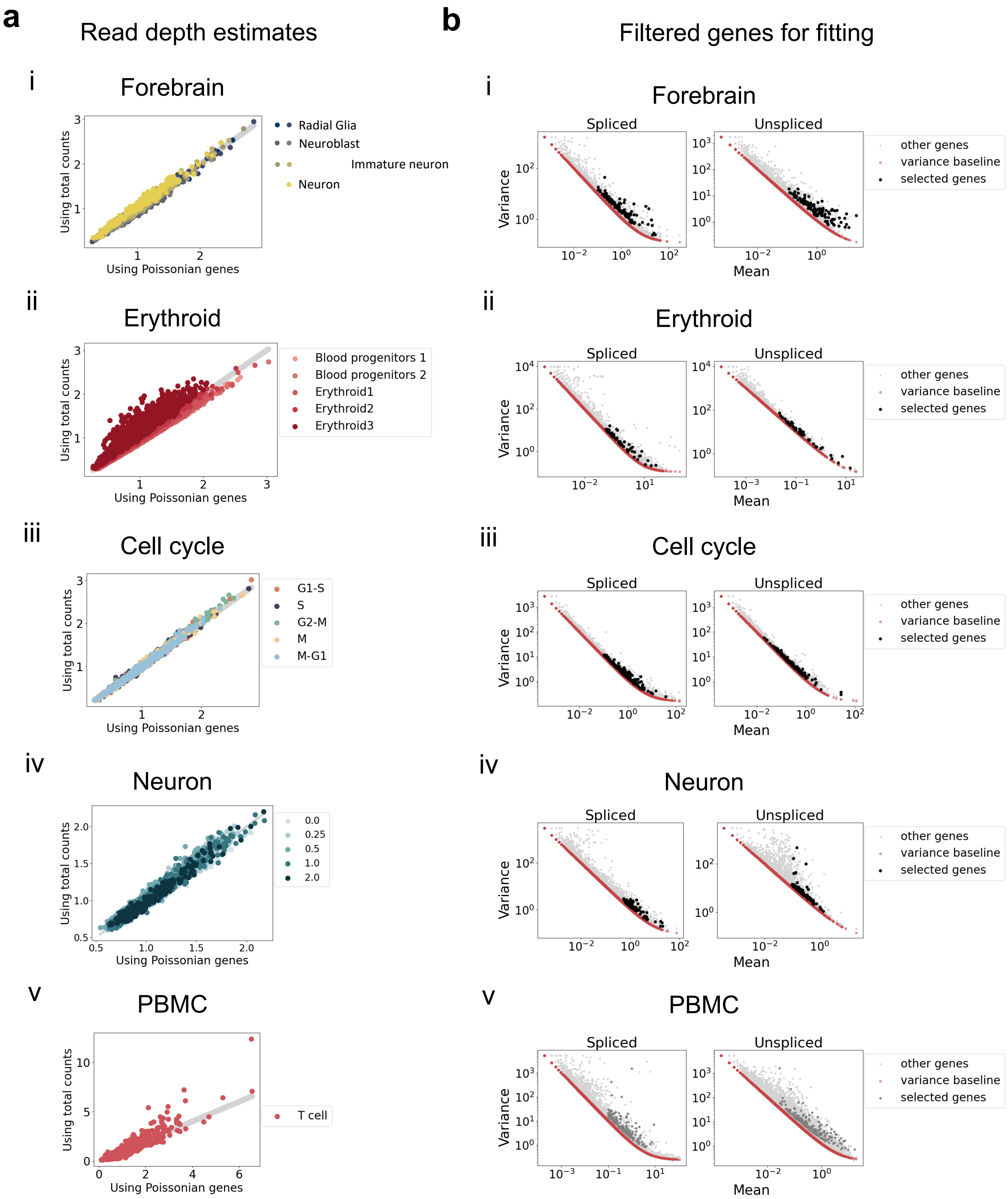


Figure S10: **Read depth estimation using normalized covariance for highly variable gene selection**. **a**) $CV^2$-mean relationship of total, unspliced and spliced counts of all genes. The baseline (red line) of variance is $1/\text{mean}$ + normalized covariance. Genes whose variance is within 1.2 times baseline are considered to be Poissonion genes (dark gray). Others are colored in light gray. **b**) Read depth estimation based on total counts of Poissonion genes. Cells are colored by their cell types. **c** Selected genes (black) for fitting plotted in the same $CV^2$-mean plot as in **a**).

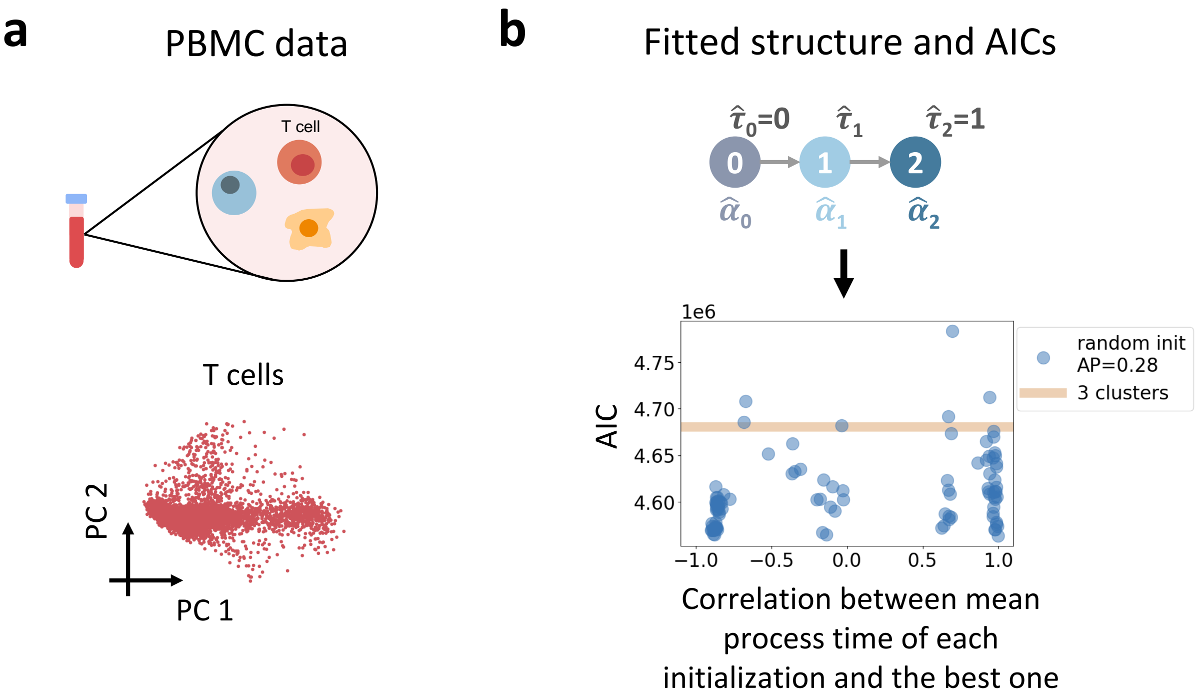


Figure S11: **Inference results for T cells from PBMC data.** **a**) Schematics of T cells from PBMC dataset and PCA plots. **b**) The fitted trajectory structure and AIC scores of 100 random initializations (blue dots) compared to those of 3 clusters (Poisson mixtures) model (yellow line). AP stands for average precision.

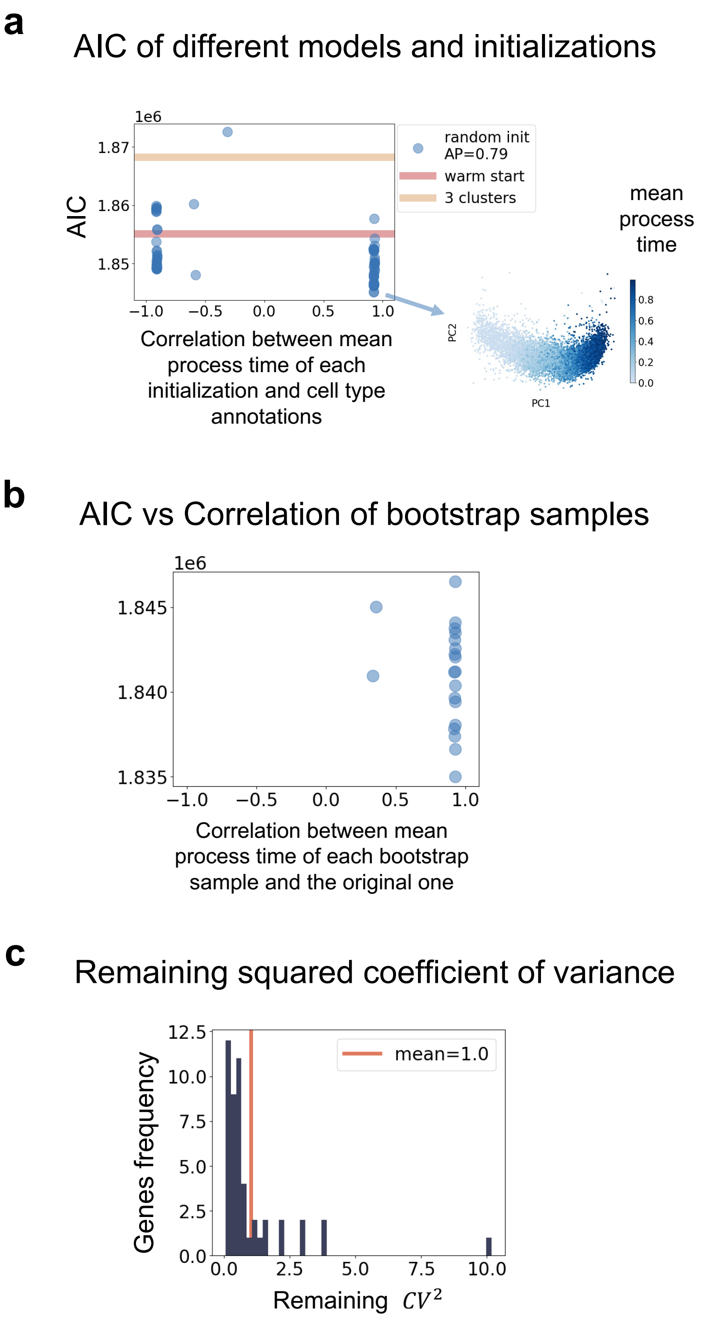


Figure S12: **Supplementary figures for Erythroid data**. **a**) AIC scores and mean process time correlations of 100 random initializations (blue dots) compared to those of warm start (red line) as well as 3 clusters (Poisson mixtures) model (yellow line). AP stands for average precision. Mean process time of the initialization with lowest AIC is indicated in blue on the same PCA plot as in **a**. **b**) AIC scores and mean process time correlations of 100 bootstrap samples. The x axis is the Pearson's correlation between the mean process time of each bootstrap and the those of original data, i.e., the plotted one in **a**. **c**) Distribution of remaining squared coefficient of variance of 49 genes used in the fitting. Remaining squared coefficient of variance is calculated by dividing the remaining unexplained variance by mean squared.

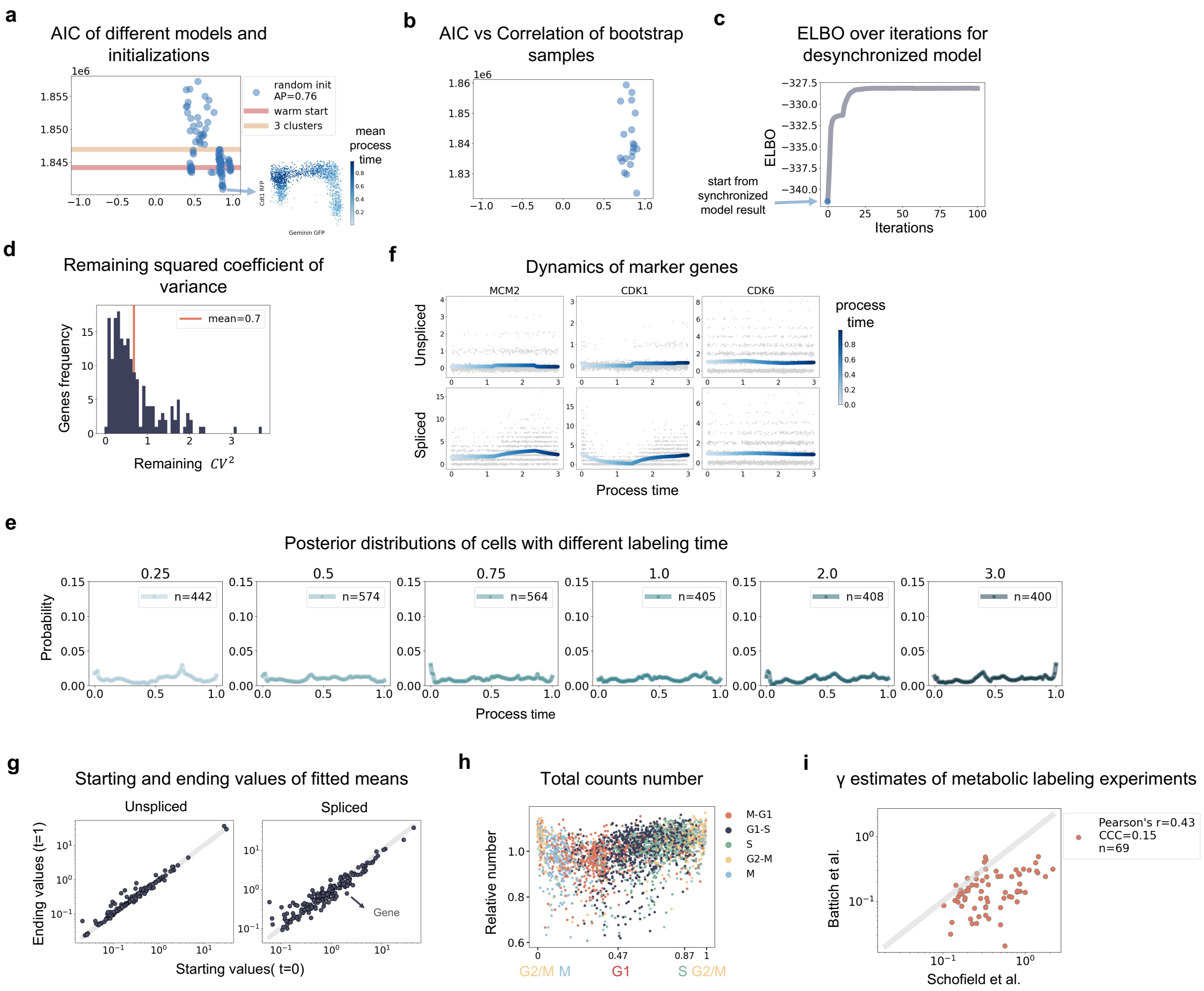


Figure S13: **Supplementary figures for cell cycle data**. **a**) AIC scores and mean process time correlations of 100 random initializations (blue dots) compared to those of warm start (red line) as well as 3 clusters (Poisson mixtures) model (yellow line). AP stands for average precision. Mean process time of the initialization with lowest AIC is indicated in blue on the same PCA plot as in **a**). **b**) AIC scores and mean process time correlations of 100 bootstrap samples. The x axis is the Pearson's correlation between the mean process time of each bootstrap and the those of original data, i.e., the plotted one in **a**. **c**) ELBO scores over iterations for desynchronized model. The fitting started with the best random initializations result of synchronized model. **d**) Distribution of remaining squared coefficient of variance of 182 genes used in the fitting. Remaining squared coefficient of variance is calculated by dividing the remaining unexplained variance by mean squared. **e**) Averaged posterior distribution across cells with different labeling times. n is the number of cells. **f**) Dynamics of three marker genes. The blue curve is the fitted mean of product Poisson distributions of unspliced and spliced counts over process time, and its darkness corresponds to the value of process time. Cells' raw counts (gray) are plotted against their corresponding process times. **g**) Starting and ending values of fitted mean of Poisson distributions. **h**) Total counts over process time of cells colored by cell type annotations. **i** Comparison of $\gamma$ estimates from two metabolic RNA labeling papers for 84 selected genes. Estimates of 67 genes are available in both papers. CCC stands for concordance correlation coefficient.

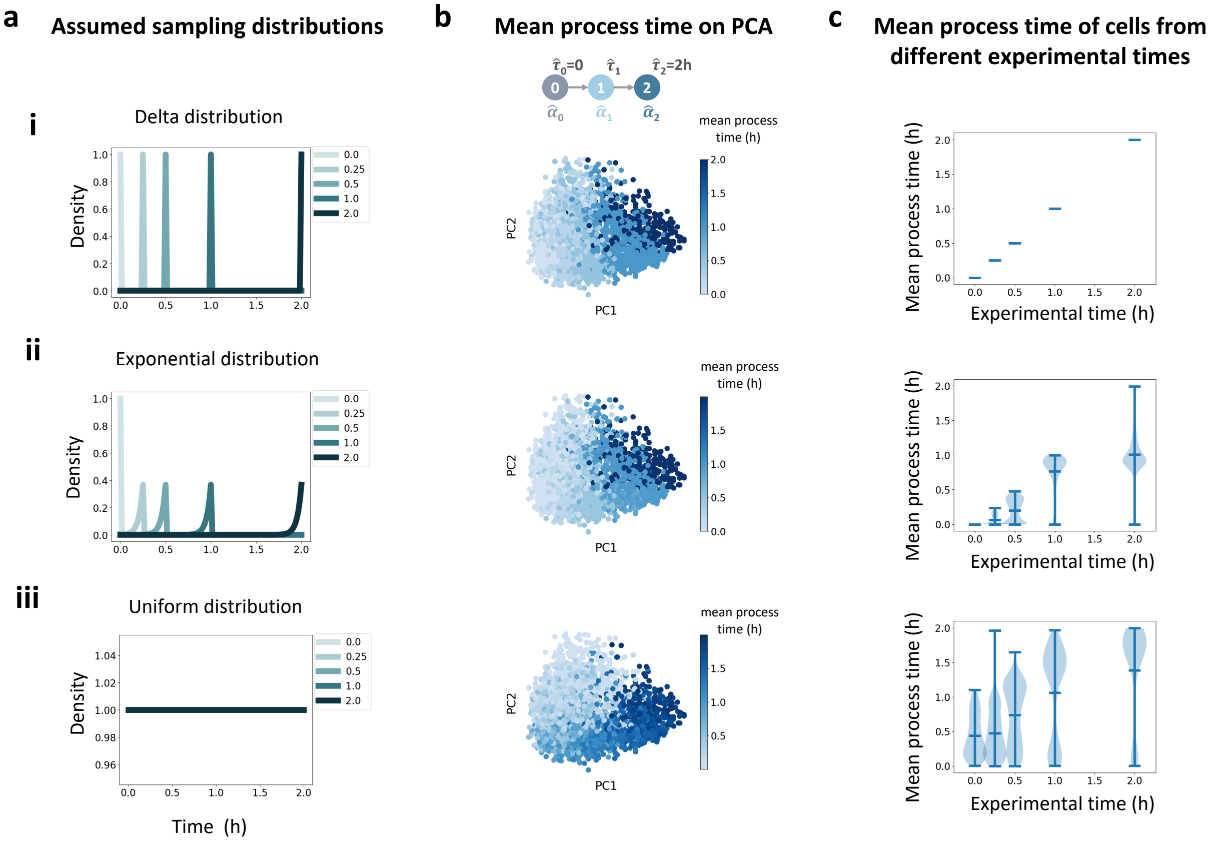


Figure S14: **Inference results of different sampling distribution assumption for Neuron data**. Fitting was warm started from delta distribution at physical time under different sampling distribution priors. **a**) The assumed sampling distribution. For **iii**, uniform distribution is assumed for cells from all time points. **b**) The fitted trajectory structure and inferred mean process time indicated in blue on the PCA plot. **c** Violin plots of mean process time of cells with different labeling times. Three blue bars represent the mean and extremes.
